# Complex genomic ancestry in southern regions and drivers of continental-level genetic diversity in the wolves of Asia

**DOI:** 10.1101/2024.09.18.613796

**Authors:** Lauren M. Hennelly, Barbara R Parreira, Ash Noble, Camilla Scharff-Olsen, M. Çisel Kemahlı Aytekin, Çağan H. Şekercioğlu, Pavel Kosintsev, Ladislav Paule, Pavel Hulva, Hans K. Stenøien, Bilal Habib, Hira Fatima, Ghulam Sarwar, Samara P. El-Haddad, Frank Hailer, Xin Sun, Nuno Filipes Gomes Martins, M Thomas P Gilbert, Mikkel-Holger S. Sinding, Benjamin N Sacks, Shyam Gopalakrishnan

## Abstract

Gray wolves (*Canis lupus*) in Asia encompass most of the species’ global genetic diversity and many endangered populations. However, a clear understanding of the evolutionary history of wolves from many parts of Asia, especially southern regions, is lacking. We used 98 whole genomes of wolves sampled across Eurasia to better resolve their evolutionary history by investigating phylogenetic and gene flow histories across the genome, and to characterize their demographic history and genetic diversity. The strongest barriers to gene flow coincided with boundaries separating the three major extant wolf lineages - Indian, Tibetan, and Holarctic. Wolves in the central Asian mountain ranges belonged to the Holarctic lineage, and despite their geographic adjacency only share minimal ancestry with the Tibetan lineage. In contrast, wolves from eastern Asia share population-wide ancestry with the Tibetan lineage, which may reflect an unsampled lineage similar, but not exactly to, the modern-day Tibetan lineage. Wolves from southwestern Asia also share population-wide ancestry with the Indian lineage, likely due to old (>6kya) admixture events. Long-term historical declines over the last 100,000 years, geographic isolation, and recent inbreeding have resulted in the Indian and Tibetan wolves having some of the lowest levels of genetic diversity and highest realized genetic loads. In contrast, adjacent populations exhibit some of the highest genetic diversity, due in part to admixture along contact zones. Our study illustrates how using multiple approaches that consider heterogenous signals across the genome can more fully resolve the historical and contemporary processes that have led to present-day species’ diversity.

## Introduction

Understanding the processes that shape genetic variation within a species is fundamental in evolutionary biology. Genetic variation is often unevenly distributed across a species’ range, and assessing where and how it varies provides insight into underlying evolutionary processes. Because multiple evolutionary processes impact a species’ genetic variation, it has long remained a methodological challenge to distinguish between these processes, such as intrapopulation and interspecies gene flow, genetic drift, and incomplete lineage sorting. The contribution of these processes also varies temporally over a species’ evolutionary history, further challenging inferences. Recent methodological advances now consider genomes as mosaics of evolutionary history whereby different genomic regions have distinct evolutionary histories (Martin et al. 2017, Excoffier et al. 2021, Harris et al. 2022). Carefully interpreting these distinct signatures across the genome has recently improved our understanding of the evolutionary histories of notoriously complex groups (Figuiro et al. 2019, Lescroart et al. 2023, Jensen et al. 2023).

Widely distributed species offer a unique opportunity to study how genetic variation is shaped by historical and contemporary environmental changes. Because they are so widespread, their populations experience different histories of isolation and interpopulation or interspecific gene flow, often in response to past and present environmental change. Applying new methodological approaches to widely distributed species can better resolve their evolutionary histories and offer additional insight into how periodic gene flow and isolation in different parts of their distribution has given rise to present-day patterns in genetic variation within species.

Gray wolves are one of the most widely distributed and best-studied large mammals, making them a promising system to explore how historical and contemporary processes shape present-day genetic variation. Due to their scientific and public appeal, they were one of the first large carnivores to be studied using genomic tools (vonHoldt et al. 2011, Stronen et al. 2013). Over the years, genomic work of wolves has grown into large-scale studies using numerous modern and ancient genomes. This has led to critical insights on recognizing the role of hybridization in speciation and impacts of the late Pleistocene on species’ persistence (Gopalakrishnan et al. 2018, Bergstrom et al. 2022). However, most such studies have been on North American and European portions of the range. Fewer genomic studies have focused on wolves in Asia, which today holds most of their phylogenetic diversity and many endangered populations.

Gray wolves are composed of three extant divergent lineages – the Indian, Tibetan, and Holarctic – all of which occur in Asia (Wang et al. 2020, Hennelly et al. 2021, Wang et al. 2022). These lineages diverged > 100,000 years ago after which the Indian and Tibetan lineages became isolated within separate glacial refugia (Wang et al. 2020, Wang et al. 2022). In contrast, wolves of the Holarctic lineage, spanning northern latitudes from northern Europe to Siberia, as well as North America, were highly connected by gene flow during the past 100,000 years (Bergstrom et al. 2022, **Figure S1**). This high gene flow resulted in genome-wide homogenization of wolves in northern Eurasia during the late Pleistocene (Bergstrom et al. 2022). While this has provided a broad picture of wolf evolutionary history in Eurasia, few studies have examined how ancestral composition and genetic diversity is distributed in wolves across Eurasia, especially in more southern regions of Asia where lineages occur in close proximity.

For example, earlier work based on from a handful of wolf genomes from southwestern Asia (defined as the region spanning Iran to the Arabian Peninsula to Türkiye) largely reflected Holarctic ancestry, with some minor ancestry from the Indian lineage, as well as from another canid species, the African wolf (*Canis lupaster*) (Gopalakrishnan et al. 2018, Hennelly et al. 2021). However, a fuller understanding of this population’s history has been hampered by a lack of geographically spread samples from across southwestern Asia. For wolves in Japan and parts of China, previous work found they contain partial ancestry similar to ancient Pleistocene wolves from ∼35-50kya, all of which fell within the Holarctic lineage based on mitochondrial DNA (Niemann et al. 2020, Madrigal et al. 2021, Segawa et al. 2022). In addition, some of their partial ancestry was found to be ancestral to late Pleistocene wolves from northern Eurasia, including ancient wolves dating back to 100 kya (Bergstrom et al. 2022). Together, this suggests various wolf populations in the southern regions of Asia may retain diversity that is ancestral to northern Eurasian wolves, and thus, were not completely homogenized by gene flow from northern wolves during the late Pleistocene (Bergstrom et al. 2022). It remains unclear what the geographical extent of ancestral diversity (defined as ancestral to Holarctic wolves in the northern latitudes) within these wolf populations is, and the spatiotemporal origin of this ancestry in their genomes. To resolve this, additional samples and analyses are needed to clarify the ancestry composition of wolves in southern regions of Asia, and what historical processes have given rise to modern-day wolves in these southern regions.

We sought to address these four gaps in our knowledge about wolves in Asia through building a genomic dataset that includes 20 newly sequenced wolves sampled from key regions and 81 previously published wolf genomes from Eurasia and North America.

Our first objective was to characterize continental-level population structure and gene flow patterns of wolves across Eurasia, with a focus on the southern regions of Asia. Our second objective was to evaluate the presence and clarify the origin of ancestral diversity (i.e. ancestral to Holarctic wolves in the northern latitudes) for wolf populations in the central Asian mountain ranges and eastern regions of Asia by incorporating recombination rate variation into phylogenetic analyses. Our third objective was similar to objective two, but evaluating ancestral diversity within the southwestern Asian wolf population using phylogenetic analyses and demographic modeling. For our fourth objective, we use our genomic dataset to characterize the genetic diversity of wolves throughout Eurasia, as well as inbreeding and genetic load.

## Results

For this study, we resequenced new whole genomes of 20 wolves from various regions in Eurasia. These regions included southwestern Asia (Lebanon, n=1; Israel, n=1, Türkiye=1), Indus plains of Pakistan (n=2), central Asian mountain ranges (Sulieman, n=2; Karakoram, n=2; Hindu Kush, n=3), the Eurasian steppe (Kazakhstan, n=3; Russia, n=3), and in Europe (Slovakia, n=1; Ukraine, n=1). We also conducted additional (i.e., deeper) sequencing of a previously sequenced Indian wolf from Central India. We compiled these newly sequenced genomes with 81 previously sequenced wolves from North America (n=3) and Eurasia (n=78), as well as dogs (n=5), and related canid species (n=12) into a dataset, which we then aligned to the domestic dog genome assembly (canFam3.1) (**Table S1**).

### Population structure and gene flow patterns of wolves in Eurasia

Principal component analysis (PCA) and inferred individual admixture proportions of 98 Eurasian wolves primarily supported the presence of three major clusters corresponding to the three major extant wolf lineages: the Indian, Tibetan, and Holarctic wolves (**Figure 1A,1B, S2**). Some wolves fell in-between these main clusters, forming clines between Tibetan and Holarctic as well as between Indian and Holarctic clusters. In both cases, these intermediate individuals came from areas nearby the corresponding contact zones between the lineages. Intermediate individuals on the Tibetan-Holarctic cline included two wolves from Tajikistan (approximately 24% Tibetan ancestry at K=6) and two wolves from Ladakh, which is a region on the southwestern edge of the Tibetan plateau. Intermediate individuals on the Indian-Holarctic cline included only wolves from Pakistan. In contrast, all 13 wolves from southwestern Asia clustered tightly nearby other Holarctic wolves and exhibited ∼15-20% Indian wolf ancestry at K=6. A few wolves, most sampled near Africa, showed basal canid ancestry, likely reflecting admixture with originating from the African wolves.

**Figure 1.**
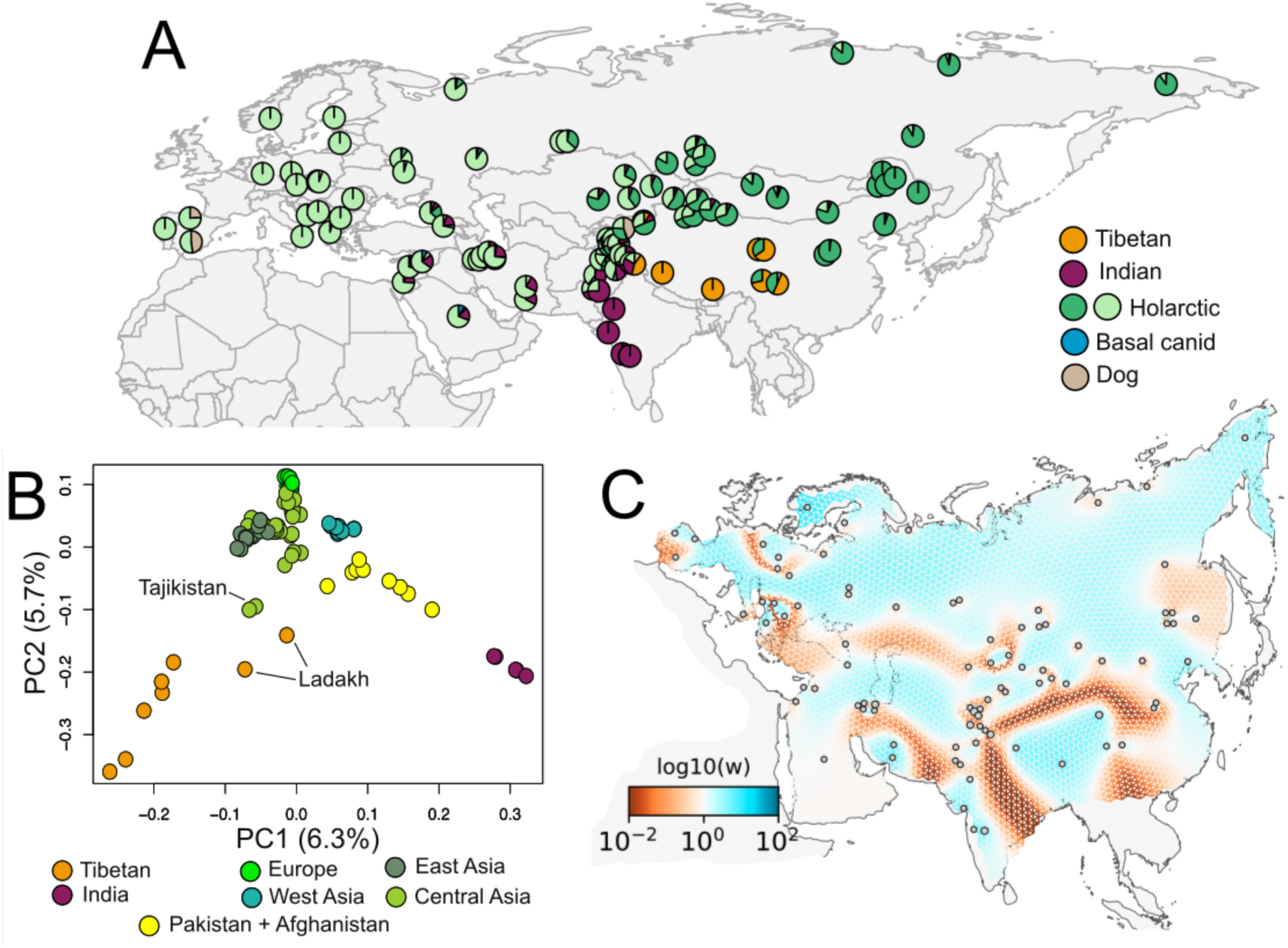
Population genetic structure genetic connectivity of wolves across Eurasia. **(A)** Sample location and individual admixture proportions estimated using genotype likelihoods of ∼300,000 autosomal SNPs at K=6 for 98 wolves across Eurasia. The individual admixture proportions were estimated using a set of 101 gray wolves, 5 dogs, and 12 genomes from various basal canid species. **(B)** Principal component analysis using genotype likelihoods of ∼10 million autosomal SNPs for 98 wolves across Eurasia. Colors indicate geographic region of each the individual wolf belongs to. **(C)** The estimated effective migration surface (EEMS) using ∼8.7 million autosomal SNPs for 97 wolves across Eurasia. Colors of the EEMS correspond to lower-than-average (red) and higher-than-average (blue) effective migration as expressed as log10(w). A lambda of 100 was used after performing cross-validation analyses.

To better characterize geographic patterns of gene flow, we inferred the estimated effective migration surface (EEMS), which indicates regions of low or high gene flow relative to an isolation-by-distance model (Marcus et al. 2021). Mirroring the PCA and admixture results, the most significant EEMS barriers to gene flow in Eurasia corresponded to divisions among the three major lineages (**Figure 1C**). The strongest barrier was along the Himalayan mountain range that separates the Indian and Tibetan lineages. A second strong barrier surrounds the Tibetan plateau, in agreement with our PCA and admixture results showing reduced gene flow between the Tibetan lineage and wolves in central and eastern Asia. The positioning of another barrier west of Pakistan was less clear, but may be artifactual due to our lack of sampling on the Eurasian steppe, such as in Turkmenistan and Uzbekistan. Otherwise, the population structure of Holarctic wolves across northern Eurasia can be modeled as isolation-by-distance (**Figure 1B,C).**

### Tibetan-like ancestry is rare in the central Asian mountains yet found population-wide in eastern Asia

To explore the evolutionary history of wolves in the central Asian mountain ranges and eastern Asia, we investigated the presence and potential source of ancestral diversity (i.e. ancestral to wolves in northern Eurasia) within these two populations. We first used D statistics to test for excess derived allele sharing with various wolf populations in Asia (Green et al. 2010). For wolves in the central Asian mountains, we observed a wide variance of allele sharing with the Tibetan lineage (**Figure 2A**). Apart from a few individuals, most wolves in Pakistan, Tajikistan and Kyrgyzstan show no statistically significant excess of derived alleles shared with the Tibetan lineage (**Figure 2A; Figure S3**). A large inter-individual variance in Tibetan derived alleles may suggest that admixture with Tibetan wolves occurs occasionally or is recent, rather than arising from long-term admixture. In contrast, we observe a consistent and significant excess of Tibetan wolf derived alleles across wolves from eastern China and Mongolia (**Figure 2A, Figure S3, S4**).

**Figure 2.**
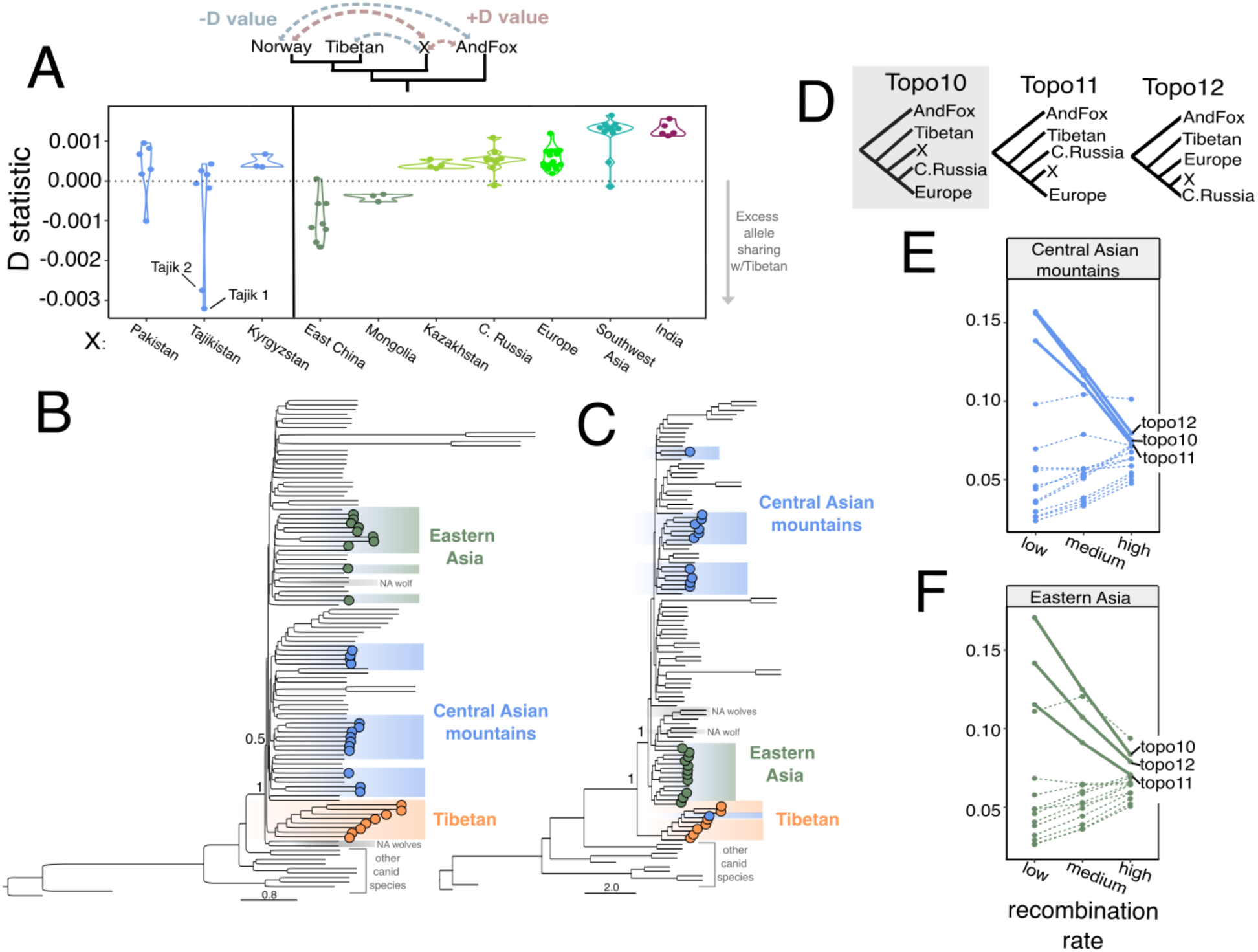
Affinity to Tibetan ancestry in wolves from central Asia and eastern Asia. **(A)** D statistic results to estimate derived allele sharing between selected wolf individuals in Asia and the Tibetan lineage. The only populations that show significant allele sharing with Tibetan wolves are wolves from Pakistan, Kyrgyzstan, Mongolia, and eastern China. **(B)** Autosomal phylogeny using 98 wolves from Eurasia, 3 North American gray wolves and 12 genomes from other canid species. The tree was inferred using ASTRAL and estimated using 1,000 regions in 20kb length that were randomly chosen across the dog reference genome. Normalized quartet score of selected nodes are shown. Colored circles indicate the individuals assigned to the Tibetan lineage (orange), Central Asian mountains (high altitude Pakistan, Tajikistan, Kyrgyzstan), North American wolves (gray), and Eastern Asia (eastern China, Mongolia). **(C)** A tree inferred using ASTRAL with only low recombination (<0.2cM/Mb) regions of the X chromosome. A total of 513K SNPs were used, which we split into regions of 10kb length to assess gene tree discordance using ASTRAL. Normalized quartet score of selected nodes are shown. Eastern Asian wolves form a well-supported clade that is basal within gray wolves, except Tibetan wolves. **(D)** The three topologies with the highest average topology weightings across the X chromosome when considering wolves from the central Asian mountains or eastern Asia as the focal population (X). The topology shaded in gray depicts a scenario where the focal wolf population is ancestral to other Holarctic wolves from Europe and Central Russia. Prevalence of 15 possible topologies in high, medium and low recombination regions on the X chromosome to assess whether wolves from **(E)** central Asian mountains and **(F)** eastern Asia show evidence of ancestry that is ancestral to northern Holarctic wolves. For both central Asian and eastern Asian wolves, three topologies (solid lines) show a negative relationship between recombination rate and topology weighting, in contrast to the 12 other topology (dotted lines). **(F)** Eastern Asian wolves have more thoroughly sorted windows when they form a lineage ancestral to other northern Holarctic populations (topology 10) in low recombination regions compared to wolves from the mountains of central Asian mountains.

It can be difficult to identify the actual source using D statistics because any ancestry more closely related to the Tibetan wolf will result in a negative D statistic. To better understand the source and historical processes that may have resulted in Tibetan-like ancestry, we inferred a set of phylogenetic trees from different regions of the genome. We first inferred a multispecies coalescent tree in ASTRAL using 5,000 regions with a length of 20kb across the autosomes using all 113 wild canid samples (Zhang et al. 2018). Tibetan wolves formed a well-supported branch basal to other Eurasian gray wolves (**Figure 2B**). The rest of the wolves form weakly supported clades, including Indian wolves which are clustered with wolves from Pakistan, and sister to those from southwestern Asia (**Figure S5**).

Because the relationships inferred by genome-wide phylogenies can be impacted by many processes, such as gene flow and incomplete lineage sorting (ILS), we leveraged how genome architecture relates to phylogenetic signal. Briefly, the X chromosome and low recombination regions are more likely to retain the historical relationship due to higher selection against introgressed ancestry and higher lineage sorting (Schumer et al. 2018, Martin et al. 2019). In particular, we took advantage of the negative relationship between recombination and historical phylogenetic relationship that has been demonstrated across multiple taxa, including wolves (Li et al. 2019, Hennelly et al. 2021, vonHoldt et al. 2021, Feng et al. 2023, Jiang et al. 2024, Monthey and Spellman 2024).

Our phylogenetic tree based on low-recombination regions (<0.2 cM/.Mb) of the X chromosome demonstrated that one of the two wolves from Tajikistan that showed the highest allele sharing with Tibetan wolves clustered within the Tibetan lineage, and all other wolves from Tajikistan and Kyrgyzstan clustered with other Holarctic wolves (**Figure 2C, Figure S6, Figure S7**). Wolves from eastern Asia formed a well-resolved clade that branched basally to the rest of the Holarctic lineage. Our phylogeny using low recombination regions (<0.2 cM/Mb) of the autosomes and the entire X chromosome recapitulates this pattern (**Figure S8, Figure S9**). In addition, topology weighting analyses show low recombination regions of the X chromosome are more resolved for topologies where wolves from eastern Asia form the basal branch within the Holarctic lineage (**Figure 2D, Figure S10**). Thus, while wolves in Tajikistan, Pakistan, and eastern Asia have similar D-values, they have contrasting topological patterns. This is consistent with wolves from Tajikistan and Pakistan having experienced recent and/or periodic gene flow from Tibetan wolves near the secondary contact zone, yet this Tibetan ancestry has not yet spread widely through the population. In contrast, wolves in eastern China and Mongolia have retained population-wide ancestry that is more ancestral than wolves in northern Eurasia, yet appears distinct from modern-day Tibetan wolves.

### Population-wide Indian ancestry within wolves of southwestern Asia

We detected complex ancestry patterns in wolves across southwestern Asia. Estimated admixture proportions and D statistics suggest that wolves in this region shared ancestry similar to the Indian lineage and another canid species, the African wolf (**Figure 1A,B**, **Figure 3A,B**). We detected nine wolves that showed significant derived allele sharing with the African wolf, which were largely confined to areas most geographically close to Egypt (Saudi Arabia, Syria, Lebanon) (**Figure S11, S12**).

**Figure 3.**
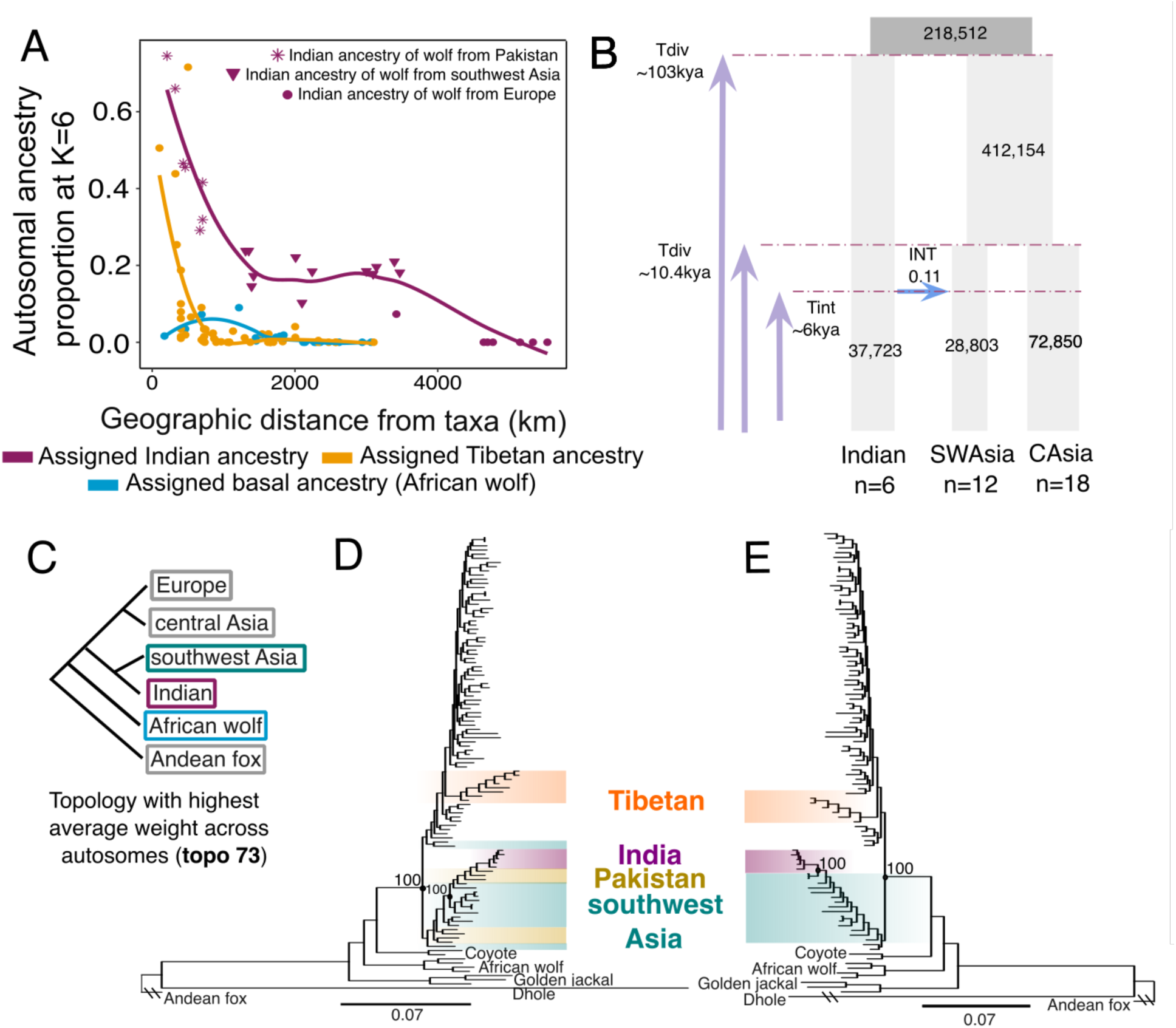
Persistence of Indian-like ancestry in wolves of southwest Asia. **(A)** Declines in autosomal ancestry proportion at K=6 assigned to the Indian lineage (purple), the Tibetan lineage (orange) and African wolf (*Canis lupaster*) (blue) for selected wolves as geographic distance increases from the distributional extent of these three taxa. Assigned Indian ancestry does not decline with geographic distance for wolves in southwest Asia (purple triangles). **(B)** The model with the highest likelihood inferred using fastsimcoal2 corresponding to an introgression event between the Indian lineage and wolves of southwest Asia. **(C)** The topology with the highest average topology weight when testing the relationship between 4 wolf populations and the African wolf, resulting in 105 possible topologies inferred using 100-SNP windows with Twisst. **(D)** A maximum likelihood phylogeny using autosomal genomic regions in which the topology weight of topology where Indian and southwest Asian wolves are sister (topology 73) was above 0.3 (∼99.99^th^ percentile; ∼40kb length total). The tree was inferred using IQ-tree where the best model was TVM+F+R5 through ModelFinder and 1000 bootstraps were used. Local posterior probability is labeled at selected nodes. **(E)** A maximum likelihood phylogeny that excludes wolves from Pakistan and Ladakh, which was inferred using autosomal genomic regions where the topology weight of topology 73 was above 0.3 (∼99.99^th^ percentile; ∼40kb length total).

Interestingly, we find ancestry shared with the Indian lineage was nearly equal in all wolves in southwestern Asia (**Figure 3A, S12, S13, S14**). There appeared to be no geographic decline in the admixture proportions of assigned Indian ancestry spanning the contact zone in Pakistan until reaching the Caucasus mountain range and from Türkiye into Europe, where no Indian wolf ancestry was detected (**Figure 3A, Figure S14**). There was also a lesser, yet significant, signal of ancestry similar to the Indian lineage on parts of the Eurasian steppe in Central Asia, such as in Kazakhstan (**Figure S14**). This geographically widespread signal contrasts what we observed with Tibetan and African wolf ancestry, which declines quickly as distance increases from their respective distributions (**Figure 3A**).

The balanced Indian wolf-like ancestry across southwestern Asia could be consistent with an admixture event between Indian and Holarctic lineages. To investigate the origin of this Indian-like ancestry, we used FASTSIMCOAL2 (Excoffier et al. 2021) to test three demographic scenarios where (A) there is no introgression event between Indian wolves and southwestern Asian wolves, (B) southwestern Asian wolves experience an introgression event with Indian wolves, and (C) southwestern Asian wolves are formed as a hybrid lineage between Indian and central Asian wolves (**Figure S15**). We obtained similar likelihood scores for the three models, suggesting no one model is strongly favored over the others (**Table S2**). However, topologies that included bifurcation (Divergence and Introgression models) had the highest likelihoods. The model that included introgression from Indian wolves to southwestern Asian wolves was preferred, with a relative likelihood of 62% (**Table S2**). Estimates suggest a relatively high introgression proportion (0.11) from India to southeast Asia, consistent with our D statistic results (**Figure S14**). We estimate the introgression occurred around 6,000 years ago, relatively close to the divergence time between southwest and central Asian populations about 10,300 years ago. Therefore, if introgression did occur, it likely happened near the time of the split between southwest and central Asian populations, explaining the similar likelihood between the divergence (A) and introgression (B) models. Importantly, both models estimate similar dates for the split between southeastern and central Asian lineages and all models date the divergence of Indian wolves from Eurasian wolves to around 100,000 years before present, providing robustness to time estimates. Additionally, both introgression and divergence models produced similar estimates for population sizes, suggesting a small effective population size (*NE*) for India and southwest Asia, whereas central Asian populations and the ancestral Asian population had a larger *NE* (**Figure S15**). Our results indicate that the Indian wolf ancestry within the genomes of southwestern Asian wolves reflects older admixture (>6kya) and/or shared demographic history rather than appearing within the past few last thousands of years.

Finally, to better assess how this Indian wolf-like ancestry relates to modern-day Indian wolves, we inferred a set of phylogenetic trees to investigate the topological positions of Indian and southwestern Asian wolves. In almost all inferred autosomal and X chromosome trees, we observe the Indian lineage and wolves from Pakistan cluster together, with almost all southwestern Asian wolves as a sister clade – all within the Holarctic clade (**Figure S5, S6, S7, S8**). The clustering of the Indian lineage with wolves in Pakistan implies they share more recent or prevalent ancestry, consistent with this region being a prominent contact zone (Hennelly et al. 2023). When excluding wolves from Pakistan and southwestern Asia, Indian wolves form the most basal branch of the gray wolf phylogeny, in line with southwestern Asian wolves being the admixed population (Wang et al. 2022, **Figure S17**). To investigate Indian-like segments within southwestern Asian wolf genomes in more detail, we identified genomic regions across the autosomes where Indian and southwestern Asian wolves were sister to each other using topology weighting with Twisst (Martin and Belleghem 2017). For this, we quantified the frequency of 105 possible topologies using 100-SNP windows across the genome for five populations with the Andean fox (*Lycalopex culpaeus*) as an outgroup (**Table S3**). The topology with the highest average weight was one where southwestern Asian wolves were sister to Indian wolves, suggesting some of their genome shares a more recent common origin rather than with other Holarctic wolves (**Figure 3C, Table S3**).

We then inferred a maximum likelihood tree using only genomic regions that are most reflective of Indian and southwestern Asian wolves being sister clades (>0.3 topology weight for the highest supported topology; **Figure 3C**). In this maximum likelihood phylogeny, we find the most basal wolf lineage consists of Indian and southwestern Asian wolves, where southwestern Asian wolves fall outside of the diversity within the modern-day Indian lineage (**Figure 3D, 3E, S18, S19**). One hypothesis for this finding is that Indian-like ancestry within southwestern Asia may derive from a once more diverse Indian lineage rather than arising directly from modern-day Indian wolves within the Indian subcontinent. Alternatively, there could be some Holarctic ancestry within these 100-SNP windows, thus placing southwestern Asian wolves outside of modern-day Indian wolf diversity. Our work emphasizes a complex evolutionary history of wolves in southwestern Asia where additional studies are necessary to fully resolve it.

### Historical and present-day impacts on genetic diversity

Our whole-genome dataset allowed us to assess patterns of genome-wide diversity of wolves across Eurasia and North America. We calculated autosomal heterozygosity using genotype likelihoods for 101 wolves from Eurasia (n=98) and North America (n=3), and found wolves in Asia encompass a wide range of some of the lowest and highest levels of genetic diversity among wolves in our dataset (**Figure 4A**). Along with the well-known depleted heterozygosity of Mexican and Iberian wolves, we found wolves in India and on the Tibetan plateau have some of the lowest documented heterozygosity in our dataset. As observed in our Pairwise Sequentially Markovian coalescent (PSMC) analysis, this remarkably low heterozygosity is in part due to their long-term population declines in the last >100 kya rather than, or in addition to, anthropogenic-related declines (Li and Durbin 2011) (**Figure 4B**). We also observe that the wolf from Shanxi, China, had a lower population size throughout the Pleistocene compared to other Holarctic wolves (**Figure 4B**). This may suggest the Shanxi wolf and other wolves in southern China were less connected to northern wolf populations, however, a greater number of higher coverage genomes are needed to rule out the effects of low coverage. Otherwise, there is a shift to elevated heterozygosity coinciding with secondary contact zones between the Holarctic lineage and the Indian and Tibetan lineages, respectively. For example, admixed wolves in lowland Pakistan have higher genome-wide heterozygosity than wolves of Central India, even though both populations are similarly endangered and declining. We find the highest heterozygosity among wolves in Asia is geographically clustered in Iran and Xinjiang of China (**Figure 4A**). These findings are concordant with our estimated inbreeding coefficients using genotype likelihoods (Hanghoj et al. 2019) (**Figure S20).**

**Figure 4.**
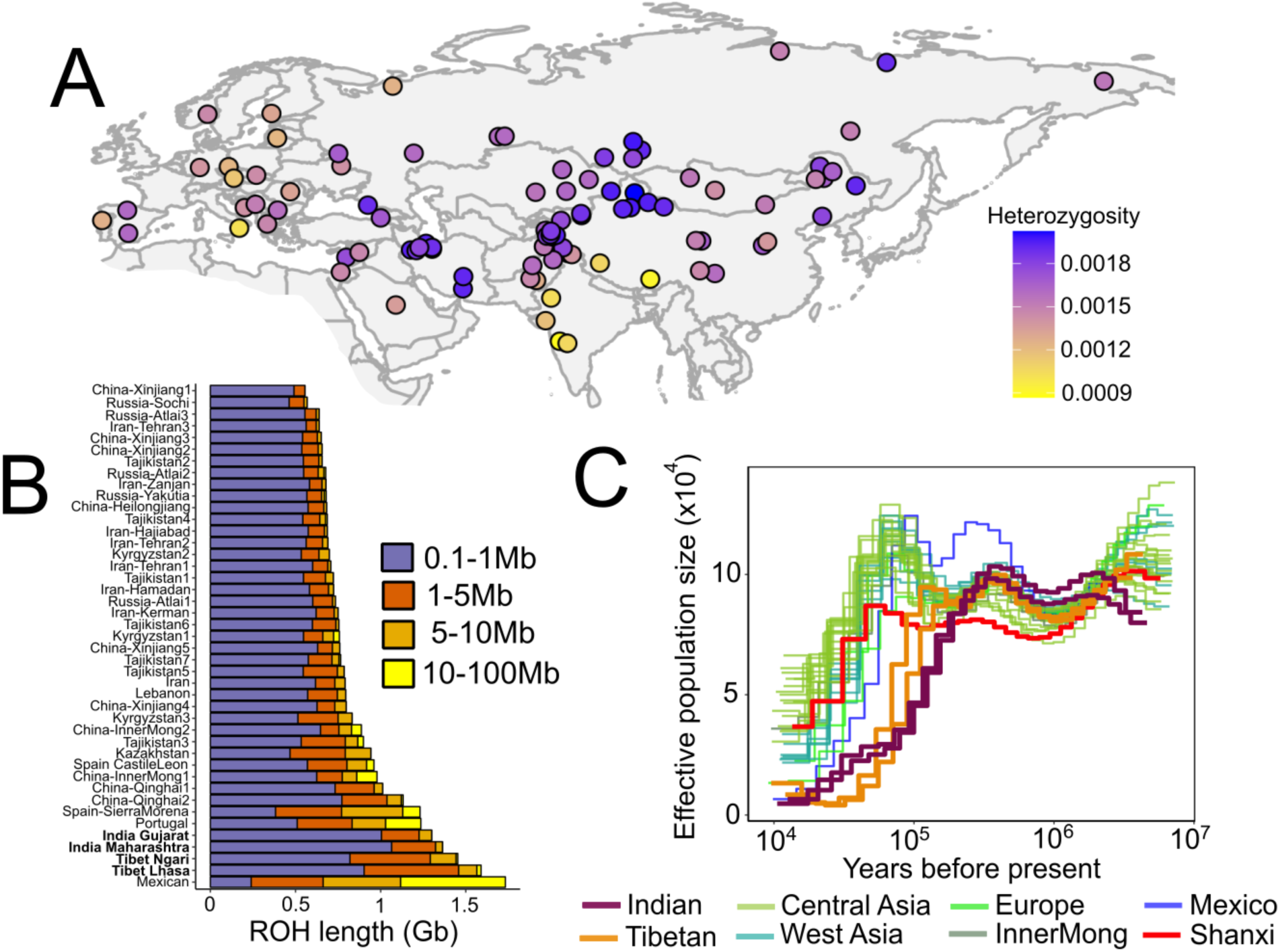
Recent and long-term demographic history of wolves in Eurasia. **(A)** Estimates of genome-wide heterozygosity inferred using ANGSD for 99 wolves from Eurasia**. (B)** Runs of homozygosity (ROH) for 43 wolves showing the sum of the length of ROH in four size ranges: 0.1-1Mb (Short ROH), 1-5Mb (Short-medium ROH), and 5- 10Mb (medium-long ROH), and 10-100Mb (long ROH). Individuals are arranged from top to bottom by the sum length of ROH with wolves most representing the Indian and Tibetan lineages bolded. **(C)** Pairwise sequentially Markovian coalescent (PSMC) analysis of the demographic histories of 33 wolves representing multiple populations in Eurasia and North America. Thicker lines indicate population belonging to the Indian and Tibetan lineages, and a single individual from Shanxi, China. We assumed a mutation rate of 4.5 x 10^-9^ and a generation time of 4.4 years.

To further disentangle what historical and present-day processes may explain patterns of genetic diversity in wolves, we inferred runs of homozygosity (ROH) for genomes with moderate coverage (>15x) using BCFtools/RoH (Narasimhan et al. 2016). The lengths of ROHs allowed additional insight into the timing of inbreeding. Combined with knowledge of the generation time and recombination rate, we estimated the timing of inbreeding, where, for example, a ROH length of 5Mb suggests inbreeding took place ∼7 generations ago (Thompson 2013). We find that Indian and Tibetan wolves show the most extensive inbreeding between 800-102 years compared to all other wolves in our study (**Figure S21**). Along with the highest number of short ROHs (0.1-1Mb), Indian and Tibetan wolves show evidence of larger ROH as well (5-10Mb), suggesting their low heterozygosity is a combination of long-term isolation over the last 200kya and anthropogenic pressures in the last hundreds of years (**Figure 4**, **Figure 3B**). We find the well documented and dramatic population declines of Mexican and Iberian wolves occurred more recently over the course of the last century, in line with previous work (**Figure S21**, Taron et al. 2021, Gomez-Sanchez et al. 2018). Likewise, we observe a few individuals in Asia possess long ROHs (5-100Mb), implying the slightly reduced genetic variation in these wolves is due to recent declines in the last 100 years (**Figure S21**). Thus, in contrast to genetic diversity patterns in Europe, our results suggest much of the variation in genetic diversity in Asia can be explained by historical processes, coinciding with glacial refugia and secondary contact zones.

Because Indian and Tibetan wolves show low genetic diversity, we explored how another component of genetic diversity – functional genetic diversity – compares to other wolves across Asia. We were specifically interested in quantifying patterns of deleterious variation that is expected to have a negative impact on fitness. Using SnpEff to estimate functional impact of mutations, we quantified the total genetic load (total number of derived alleles classified as high or moderate impact), realized load (homozygous state of derived alleles in high or moderate impact), and masked load (heterozygous state of derived alleles in high or moderate impact) for five selected wolf populations with assuming the derived alleles was the deleterious one (Cingolani et al. 2012, Bertorelle et al. 2022). Wolves from historically large population sizes – central and southwestern Asia – had higher masked and total genetic load than wolves from historically smaller populations (Indian, Tibetan, Mexican wolves) (**Figure 5A,B, S22, S23**). This is consistent with theoretical predictions that larger populations with higher genetic diversity should also possess more deleterious alleles in a heterozygous state (Mathur and DeWoody 2021). In contrast, we found that Indian, Tibetan, and Mexican wolves have a higher realized load and a lower total genetic load (**Figure 5,C**) than the other surveyed populations. A higher realized load in these small populations is expected, because genetic drift and inbreeding will result in an increased proportion of homozygous genotypes, including deleterious ones.

**Figure 5.**
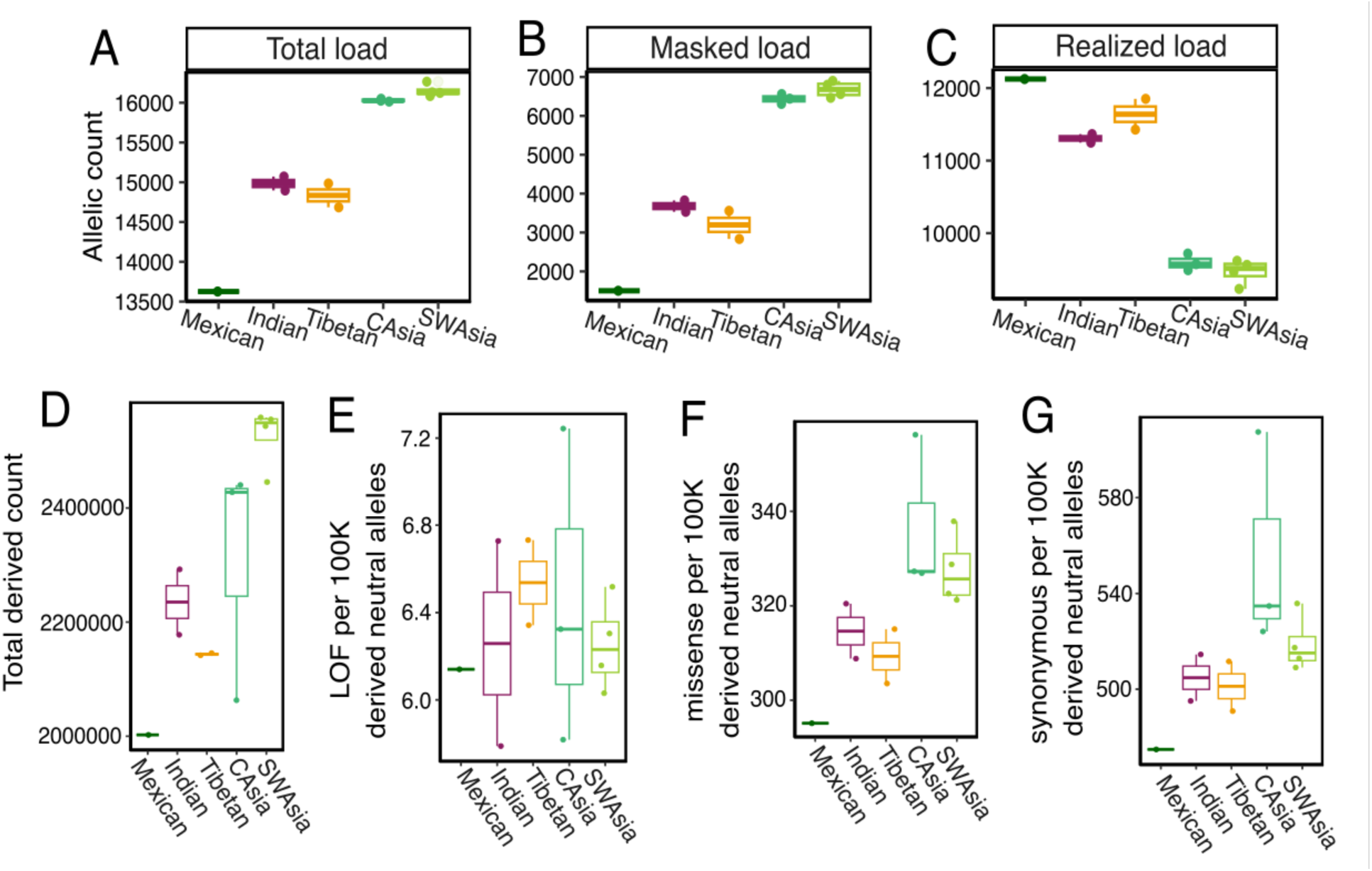
Genetic load in selected wolf individuals in Asia and North America. **(A)** Total genetic load, **(B)** masked load, and **(C)** realized load for 13 wolf individuals wihtin five populations. Wolves found in historically smaller populations (Mexican, Indian, and Tibetan) have a lower total load, lower masked load, and higher realized load. The total derived allele count of each wolf within the five populations, where generally Mexican, Indian, and Tibetan wolves show overall less derived alleles. The derived allele count of **(E)** Loss of function (LOF) mutations classified as high impact, **(F)** missense mutation classified as moderate impact, and **(G)** synonymous mutations classifed as low impact per 100,000 derived alleles found in non-coding regions, assumed to be neutral. When taking into account differences in overall number of derived count, Mexican, Indian, and Tibetan wolves show reduced number of missense and synonymous mutations, but not for loss of function mutation, compared to central Asian and southwest Asian wolves.

A lower total genetic load in the Indian and Tibetan wolves is intriguing, as this may reflect a loss of derived variants due to stronger genetic drift in historically small populations and/or purging (Dussex et al. 2023). Purging predominantly occurs when recessive deleterious variation is exposed to purifying selection in a homozygous state. To look for explore evidence of stronger genetic drift versus purging, we asked whether deleterious alleles are lost at a higher frequency than the background neutral alleles (mutations in non-coding regions) for each individual wolf. In other words, does the fact that derived alleles are categorized as deleterious result in more frequent loss compare to neutral derived alleles within an individual, as expected for purging via purifying selection? Or are deleterious derived alleles lost at the same frequency as neutral derived alleles, suggesting the loss in deleterious derived alleles found in our total genetic load results could be due to genetic drift? We assessed the prevalence of sites associated with four impact categories: high impact (loss of function, LOF), moderate impact (missense), low impact (synonymous), and modifier (non-coding; here assumed to be selectively neutral). To account for differences in total derived alleles found in wolf individuals, we divided the total number of LOF, missense and synonymous derived mutations by the total number of neutral derived alleles assumed to be under neutral processes (mutations in non-coding regions) for each individual.

We found Indian, Tibetan, and Mexican wolves have proportionally fewer missense and synonymous mutations per 100,000 derived neutral mutations compared to wolves in central and southwest Asia. A reduced frequency of synonymous mutations may reflect stronger genetic drift associated with a smaller long-term population size rather than purifying selection via purging, as synonymous mutations are assumed to be nearly neutral (**Figure 5F,G**). We emphasize that it is difficult to differentiate evidence of genetic drift and purging, and further work is needed to verify the functional impacts of our missense and synonymous mutations. For high impact mutations, we observe no clear pattern among individuals (**Figure 5E**). One reason for this could be incorrectly annotated LOF mutations in the short-read assembled dog reference, as suggested by Smeds et al. (2023). In addition, among the ones that may be correct, there are so few LOF sites across the genome (n=145 on average in our dataset) and being such a small number, their frequency may be more strongly influenced by genetic drift, as seen with killer whales (Kardos et al. 2023). Overall, we suggest caution when interpreting our genetic load results. Further work is needed to assess the accuracy of annotations from a short-read assembled genome, functional impacts of mutations, and to increase the number of individuals included in each population.

## Discussion

In this study we conducted a comprehensive whole genome analysis on population structure, ancestral composition, and genetic diversity of wolves in Asia. We find that the strongest barriers to gene flow in Eurasia coincides with the Indian, Tibetan, and Holarctic lineage distributions. We show that Holarctic wolf populations found in southern regions of Asia possess genetic variation ancestral to wolves in Northern Asia and that it is similar in the degree of its antiquity to that of the Tibetan and Indian lineage. Despite evidence of recurrent gene flow across northern Eurasia during the late Pleistocene, it appears this process of homogenization did not completely reach areas in southern regions of Asia (Bergstrom et al. 2022). Lastly, for wolves in Asia we find genetic diversity largely corresponds to historical processes, such as long-term isolation within glacial refugia, contact zones, and long-standing environmental differences, followed by anthropogenic processes.

Asia is the biogeographic cradle for gray wolf evolution, so it is not too surprising that Asia harbors ancestral diversity in its southern regions. Previous work has also shown that southern regions contain deeper local ancestry than both modern and some Pleistocene wolf genomes from Northern Eurasia. Wolves from southwestern Asia, eastern parts of China, and Japan contain ancestry that is more ancestral than Siberian wolves living during the Pleistocene from 12 kya and even 100 kya (Bergstrom et al. 2022). In addition, a single wolf from Shanxi of eastern China had significant allele sharing with ancient Siberian wolves from 50kya, 48kya, and 33kya (Ramos-Madrigal 2021). Recent work has emphasized the importance of carefully interpreting Introgression analyses and to consider the possibility of “ghost lineages”, that is extinct or unsampled lineages (Tricou et al. 2022). For eastern Asian wolves, we detected significant introgression with the Tibetan lineage as P3, but not with the older diverged Indian lineage or African wolf as P3. While this suggests the origin of this partial ancestry within eastern Asian wolves is similar to modern-day Tibetan lineage, the degree to which the partial ancestry is similar to the Tibetan lineage remains unclear. Our phylogenetic trees provide some clarity. None of our trees nor most frequent topologies using Twisst show eastern Asian wolves clustering within or sister to the Tibetan lineage. Instead, low recombination regions on the X chromosome show they form their own lineage that diverges before the rest of the Holarctic wolves, yet after the Tibetan lineage. Thus, it is possible that wolves in southern China may contain ancestry of an unsampled wolf lineage that was present in southern China and was distinct from the Tibetan lineage.

Consistent with this, previous work shows historical genomes of wolves in Southern China and Tibetan wolves occupy opposite ends of the PC2 axis, suggesting distinct partial ancestries (Wang et al. 2019). Fossils of wolves have been present across most regions of China over the last 100,000 years, including 12,000 year old remains from the Southern province of Jiangxi (Wang 2016). During this time, much of southern China was drier with mixed broadleaf forests occupying where monsoon rain forests are now (Pinxian and Xiangjun 1994). Thus, it is plausible that ancestry that was distinct from the Tibetan lineage could have existed within a glacial refugium in Southern China, and has not been completely replaced by expansion waves from northern Eurasian wolves. An interesting future avenue of research is whether partial ancestry of extinct Honshu wolves may also be from this glacial refugia, and thus share a common origin with wolves of southern China. Incorporating ancient genomes from eastern China and the newly sequenced 35 kya Japanese wolf genome in future analyses would be needed to place this ancestry into a spatiotemporal context during the Pleistocene.

An unexpected result in our study was uncovering population-wide ancestral diversity present across southwestern Asia. Our analyses point towards the widespread ancestral component is likely related to the Indian lineage, rather than from the African wolf. This fits well with previous work suggesting part of the deep ancestry within southwestern Asia is at least 100,000 years old, which is before the estimated divergence time of the Indian lineage (Bergstrom et al. 2022, Wang et al. 2022). When and how the present-day population structure formed in southwestern Asia remains unclear, but our study offers some insights. Combined with the fact there have been no Indian mitogenomes detected outside of the Indian subcontinent, our demographic history results from fastsimcoal2 and PSMC finds no strong evidence that the Indian-like ancestry within southwestern Asia is due to recent (pre-6kya) gene flow from the Indian lineage. In addition, it appears that the X chromosome, including low recombination (<0.2 cM/Mb) regions, of southwestern Asian wolves are more similar to the Holarctic lineage. Along with our PSMC results, the major genomic background of southwestern Asian wolves does seem to be that of the broader Holarctic lineage, possibly tracing back to gene flow from northern latitudes. At the same time, we find the Indian-like genomic regions across the autosomes that do not fit into the modern-day diversity of the Indian lineage. These could be remnants of a once more diverse Indian lineage, or, despite our efforts, these genomic regions could contain some Holarctic ancestry that places southwestern Asian wolves nearby but not within Indian wolves. A more thorough analysis that uses multiple local ancestry approaches to classify Indian-like ancestry would be insightful, as the length distribution and its phylogenetic relationships can be more thoroughly assessed. More broadly, ancient remains of wolves have been documented in southwest Asia throughout the late Pleistocene (Kurten 1965, Dayan et al. 1992, Mashkour et al. 2008, Wang et al. 2016). Ancient genomes from these wolf remains in southwestern Asia can directly test hypotheses on whether there were shifts in ancestral composition over the late Pleistocene and if the Indian lineage was once more widespread and genetically diverse.

Together, our work uses a combination of methodological approaches to better resolve the historical and contemporary processes that shape genetic variation within a wide ranging species. For many species during the late Pleistocene, their populations underwent distinct evolutionary trajectories of isolation and expansion in response to shifting climates. The process in which populations don’t or do come together, and how often, is a key force in speciation and evolution. Our study’s findings of divergent ancestry persisting in southern regions may be common pattern in widespread species, where red foxes (*Vulpes vulpes*), Eurasian otters (*Lutra lutra*), and brown bears (*Ursus arctos*) all show evidence of cryptic divergent ancestry in southern regions of Asia (Statham et al. 2014, Jong et al. 2023, du Plessis et al. 2023). While our study addresses gaps in our knowledge on wolves, many questions remain to be answered. For example, why some wolf populations seem to have been more susceptible to gene flow from northern populations (southwestern Asia, eastern Asia) during the late Pleistocene, while others show little gene flow and prominent contact zones (Indian and Tibetan) is an open question. With so many ancient and modern genomes now available, wolves are poised to serve as a valuable model to study how intra-population gene flow, interspecies hybridization, and isolation during the late Pleistocene has contributed to present-day species’ diversity.

### Conservation Implications

Our study has several implications for the conservation of various wolf populations in Asia. For taxonomy, our study supports previous calls that a taxonomic revision is necessary for wolves in Asia (Alvares et al. 2019, Werhahn et al. 2022, Krofel et al. 2022). While wolves from India to Türkiye are collectively described as *Canis lupus pallipes*, our study emphasizes past work that the ancestral Indian lineage is confined to India and parts of Pakistan (Hennelly et al. 2021, Hennelly et al. 2023). For the wolves in southwestern Asia that are considered “*Canis lupus pallipes*”, we find this distinct population may be geographically limited by the Bosphorus strait in Türkiye and the Caucasus mountain range (Figure 1AB, Figure 2A). Further work is necessary to understand the population genetic structure in the arid areas of southern Central Asia, such as the Karakum desert in Turkmenistan and Kyzylkum desert in Uzbekistan. We suggest the morphological and genomic distinctiveness of wolves in southwest Asia warrants subspecies status, in line with recent recommendations (Werhahn et al. 2022). For Indian and Tibetan wolves, the presence of gene flow barriers and likely reproductive barriers coinciding with lineage boundaries further supports the recognition of subspecies or species status (Werhahn et al. 2022).

Our study underscores the importance of robustly describing wolves in parts of China. Wolves have been documented as South as Guangxi in China, a province that borders Vietnam (Wang et al. 2016, **Figure S1**). In 1936, wolves were reported to exist in parts of Central and South China, but were already less common compared to Northern China (Sowerby 1936). A report in 2003 concluded wolves have largely been extirpated from regions of Guangxi that once occurred there in the 1990s, but they were possibly still present in Nanling National Nature Reserve (Fellowes et al. 2003). In response to their decline in the country, the wolf was recently added in 2021 as a protected species under Class II of the List of Wildlife under Special State Protection of China. To the authors’ knowledge, we are not aware of published estimates of the population size and status, or a morphological description, of wolves in Southern parts of China. Given their evolutionary distinctiveness, our work highlights the need for more research on their distribution, genetics, morphology, and population status to conserve what populations remain.

Overall, our study emphasizes that southern regions of Asia hold most of the ancestral genetic diversity within gray wolves, and are also the most endangered. The Indian lineage numbers at ∼2,700 individuals in India and the Tibetan lineage only ∼3,000 individuals (Jhala et al. 2022, Werhahn et al. 2023). There are less than ∼750 wolves left on the Arabian peninsula, a genomically unique but understudied population where only one genome has been sequenced (Bonsen et al. 2024). It is possible that wolves in Southern China are already extinct or very close to it. Wolves in these regions are impacted by numerous human-induced threats, especially for the remaining populations in Southern China and India. Our study highlights the need to prioritize these populations for conservation to preserve the full spectrum of extant wolf diversity.

## Methods

This study includes 20 new whole genomes from wolves from Pakistan (n=8), Israel (n=1), Lebanon (n=1), Afghanistan (n=1), Türkiye (n=1), Ukraine (n=1), Slovakia (n=1), Kazakhstan (n=3), and Russia (n=3). For all wolf samples from Pakistan and Israel, we extracted DNA from these samples using DNAeasy Blood and Tissue Kit (Qiagen) following the manufacturer’s protocol. For the rest of the wolf samples, they were extracted using Thermo Scientific KingFisher instrument following manufacturer’s protocol. The Indian wolf from Maharashtra originated from a previous study and was resequenced to a deeper depth following protocols in Hennelly et al. 2021. For the gray wolves from Pakistan and Israel, libraries were sequenced on an Illumina NovaSeq 6000 S4 flowcell and paired end at 150-bp (Illumina). For the rest of the newly sequenced wolf samples, they were sent to BGI Copenhagen for library build and sequenced on ⅛ lane each on DNBSEQ at paired end 150. We constructed a dataset comprising an additional 105 previous published canid genomes, resulting in a total of 118 canid genomes in the study (**Table S1**).

### Alignment, Variant Calling, and Filtering

We used the Paleomix v.1.2.13.2. pipeline to align our raw reads to the dog reference genome Canfam3.1 (Schubert et al. 2014). Specifically, we used the BWA mem algorithm and used the GATK indel realigner to generate an indel-realigned BAM file for each sample. Variant calling was performed using the Genome Analysis Toolkit (GATK) version 4.2.5.0 (McKenna et al. 2010). First, we used GATK Haplotype Caller to perform variant calling on each sample, then used the resulting GVCF files to perform joint genotyping with all canid individuals combined using GenomicsDBImport and GenotypeGVCF in GATK. We obtained a set of high quality single nucleotide polymorphisms (SNPs) using the following filtering steps in GATK: QD < 2.0 || SOR > 3.0 || FS > 60.0 || MQ < 40.0 || MQRankSum < -12.5 || ReadPosRankSum < -8.0. Sites with a mean depth of > 1800 for all individuals were removed to exclude paralogues from our dataset. Lastly, we removed indels from our dataset using SelectVariants in GATK (flag -select-type SNP). We excluded genomes that were derived from museum samples in the genotype calling, which was only the wolf from Afghanistan.

### Principal Component Analysis and Individual Admixture Proportions

To assess genome-wide genetic structure of the autosomes, we conducted a principal component analysis (PCA) using PCangsd (Meisner and Albrecthtsen 2018) with our total set of Eurasian gray wolf genomes. We inferred the PCA using genotype likelihoods from ANGSD, in which we used the SAMtools model (-GL 1) and filtered the dataset by including only properly paired reads (-only_proper_paired 1) from autosomal bam files, excluded reads with excess of mismatches (-C 50) or mapping quality lower than 20 (-minMapQ20), removed transitions (-noTrans 1), and keep bases with a sequence quality above 20 (-minQ 20) (Korneliussen et al. 2014). We also retained SNPs that were present in at least 90% of individuals present (>90 out of xx total; - minInd 90). We obtained the covariance matrix from PCAngsd and used the R function *eigen* to infer the PCA.

To estimate individual ancestry proportions, we inferred genotype likelihoods using ANGSD and ran NGSadmix (Skotte et al. 2013) using K=2 to K-9 genetic clusters. We used the canids within the genus *Canis* in this analysis, which included an additional 3 North American gray wolves, 3 golden jackals, 2 coyotes, 4 African wolves, 1 Ethiopian wolf, 5 dogs (**Table S1**). We used the same criteria in ANGSD as to infer the PCA, however, we retained SNPs that were present in at least 90% of individuals. This resulted in 578,343 SNPs after filtering.

### Assess barriers to gene flow for wolves across Eurasia

We inferred the estimated effective migration surface (EEMS) to assess spatial variation in rates of gene flow among gray wolf populations in Eurasia (FEEMS; Marcus et al. 2012). The FEEMS approach utilizes a stepping-stone model to calculate genetic dissimilarities between individuals based on spatial and genetic data. These genetic dissimilarities can be visualized to illustrate departures from strict isolation by distance, thereby detecting regions of high (i.e. corridors) and low (i.e. barriers) to gene flow. For the FEEMS, we only used our dataset of 95 Eurasian wolves, where we filtered the dataset in which we kept sits with a minimum allele count of 3 (--mac 3), all of individuals represented at a site (--geno 0), and pruned sites in high linkage disequilibrium by removing each SNP with a r^2^ value of greater than 0.5 (--indep-pairwise 50 10 0.5). This resulted in ∼8.7 million autosomal SNPs across our Eurasian wolves. For gathering the geographic data for each wolf sample, we had some gray wolf samples that only had location information as the country or province within a country.

In these cases, we took the center coordinate within that country or province.

To run FEEMs, we first constructed a dense spatial grid using an edge width of 5 that covered the entire Eurasian continent. We initially performed a random initialization over the map to fix the estimate of the residual variance. We selected the tuning parameter, lambda, using leave-one-out cross-validation using 20 folds where the lambda with the lowest cross-validation was selected. Our cross-validation test found a lambda value of 100 to be the optimal value of lambda (**Figure S24**).

### Phylogenomic relationships

We constructed the autosomal and X chromosome phylogeny that included all wild canid species except dogs and the Ethiopian wolf (**Figure 2B, S5, S6**). We filtered the autosomal and X chromosome VCF to exclude indels, keep sites with at least minimum quality of 30 (--minQ30), keep only biallelic sites, and kept sites that had at least 90% individuals called (-max-missing 0.9). We then converted each vcf file into the phylip format using the script vcf2phylip (Ortiz et al. 2019). We randomly selected 500 and 1,000 regions with a length of 20 kb across the autosomes and X chromosome, respectively. We inferred a maximum-likelihood phylogenetic tree for each 20kb region using IQ-Tree 1.6.12 where we estimated the best model using ModelFinder and used 1,000 ultra-fast bootstraps to infer each tree (Kalyaanamoorthy et al. 2017, Nguyen et al. 2014). The inferred phylogenetic trees were used as input into ASTRAL 5.7.8, which inferred a species tree from a set of gene trees while taking into account gene tree discordance (Zhang et al. 2018).

In addition, we inferred phylogenetic trees of the X chromosome and autosomes using only the low recombination regions across the genome. For the X chromosome, we inferred five different runs containing different sets of canid individuals. These runs were of: (1) including all wild canid species except dogs and the Ethiopian wolf, (2) all wild canid species and wolf individuals excluding dogs, the Ethiopian wolf, wolves from Pakistan, and wolves from Ladakh (3) all wild canid species, and wolf individuals excluding dogs, the Ethiopian wolf, wolves from Pakistan and Ladakh, Indian wolves, (4) all wild canid species and wolf individuals excluding dogs, the Ethiopian wolf, wolves from Pakistan, and wolves from southwestern Asia, and (5) all wild canid species and wolf individuals excluding dogs, the Ethiopian wolf, wolves from Pakistan, and two wolves from that clustered with Indian wolves (wolf from Zanjan and Hamadan in Iran). All VCF files were filtered for sites with a minimum quality of 30 (-minQ 30), only biallelic sites, and at least 90% individuals present at each site (--max-missing 0.9). We downloaded the recombination map of the domestic dog from Auton et al. 2013 and used the recombination rate (cM/Mb) in our analyses. We averaged the recombination rates within 100-SNP windows across the autosomes and X chromosome, as generated in a previous study (Hennelly et al. 2021). We selected genomic regions that had an average recombination rate of below 0.2 cM/Mb, and used these positions to create a bed file, which we then extracted these genomic regions from our VCF.

We inferred low recombination phylogenies for each of the five runs with different samples with IQ-Tree 1.6.12 (**Figure S6, Figure S16, Figure S17A,B,C**). We converted the VCF containing all low recombination regions to a phylip file format using vcf2phylip.py (https://github.com/edgardomortiz/vcf2phylip). We then ran IQ-Tree 1.6.12 with estimating the best model using ModelFinder and used 1,000 ultra-fast bootstraps to infer each tree (Kalyaanamoorthy et al. 2017, Nguyen et al. 2014). Second, we inferred low recombination autosomal and X chromosome phylogenies using ASTRAL 5.7.8 (**Figure 2C, Figure S7, Figure S8**). We split each SNP dataset into 10 kb regions, and converted each of these regions to a phylip file (Zhang et al. 2018). We then inferred a tree for each 10 kb region using IQ-Tree 1.6.12 with estimating the best model using ModelFinder and used 1,000 ultra-fast bootstraps.

### Assessing genome-wide gene admixture

To test for gene flow among various canid lineages and species, we performed D-statistic analyses using AdmixTools (Patterson et al. 2012). We filtered our dataset to exclude missing data (--max-missing 1), kept biallelic sites, and a minimum quality filter of 30 (--minQ30), along with the filters we applied using GATK. We converted plink files into eigenstrat format using the convertf script within ADMIXTOOLS. We calculated the D-statistic and Z-score for modern populations for three different topologies in the format (((P1, P2),P3),P4): (((Norwegian wolf MW005, X wolf); Indian wolf BH123); Andean fox), (((Norwegian wolf MW005, X wolf); Tibetan wolf TI32); Andean fox), (((Norwegian wolf MW005, X wolf); African wolf); Andean fox). The D-statistic value will be negative for an excess of shared derived alleles between P2 and P3, and/or P1 and P4. Alternatively, the D-statistic will be positive for an excess of shared derived variants between P2 and P4, or P3 and P1. For African wolves, we used multiple individuals as a population for P3, which consisted of all four of our African wolf samples from Kenya, Ethiopia, Morocco, and Algeria.

We also quantified the frequency of different topologies in genomic windows across the genome using Twisst (Martin and Van Belleghem 2017). To prepare the dataset for the Twisst analysis, we first phased the filtered dataset that contained no indels, biallelic sites with a minimum quality of 30 and that were present in at least 90% of individuals using Shapeit v2.r904 (Delaneau et al. 2013). We used the dog genetic map from Auton et al. 2013 as input and a window size of 0.5 Mb. We converted the phased VCF to a geno file using the parseVCF.py script within https://github.com/simonhmartin/genomics_general. We then estimated phylogenies in 100-site windows using phyml with a General Time Reversible (GTR) substitution model with phyml_sliding_windows.py within https://github.com/simonhmartin/genomics_general. We summarized the relative prevalence of topologies using the twisst.py script. The set of topologies we summarized consisted of four population groups with Andean fox as an outgroup focused on wolves in Central Asia and Eastern Asia, and five population groups with Andean fox as an outgroup focused on southwestern Asian wolves. These resulting in 15 possible topologies for the four population groups, and 105 possible topologies using five population groups. We also quantified the frequency in which topologies occurred in windows of low (<0.2cM/Mb) recombination rate regions. To do this, we partitioned our our averaged recombination rates within 100-SNP windows inferred in Hennelly et al. 2021 across the autosomes and X chromosome datasets.

Finally, we inferred phylogenetic trees using genomic regions within wolves of southwestern Asia that were designated to be Indian ancestry from our twisst results. First, we normalized the Twisst results for all 105 topologies and calculated the mean for each topology. The topology with the highest mean of 0.33 was topology 73 where Indian and southwestern Asian wolves form sister clades as the most well-resolved topology across the autosomes (**Table S3**). We extracted genomic regions that had a topology weight of over 0.2 and 0.3 for topology 73 from a VCF containing all autosomal SNPs using bedtools. This resulted in 299,560 SNPs and 40,887 SNPs found within genomic regions that had over 0.2 and 0.3 topology weight for topology 73, respectively. We converted two VCF datasets, one with genomic regions that had over 0.2 and 0.3 topology weight for topology 73, to a phylip file using vcf2phylip.py (https://github.com/edgardomortiz/vcf2phylip). We then ran IQ-Tree 1.6.12 with estimating the best model using ModelFinder and used 1,000 ultra-fast bootstraps (Kalyaanamoorthy et al. 2017, Nguyen et al. 2014) (**Figure 3D,E, Figure S18, Figure S19**).

### Demographic Modeling with fastsimcoal2

In order to better understand the origin of southwestern Asian wolves, we compared three alternative demographic models and inferred demographic parameters under the FASTSIMCOAL2 (v2.7 Excoffier et al 2021), a composite-likelihood method that estimates historical introgression, divergence times and population topologies from the site frequency spectrum using coalescent simulations.

We compared three demographic models (**Figure S15**) to better assess the demographic history of southwestern Asian wolves. The included three populations representing India, southwestern Asia and central Asia samples, and were defined as (1) a divergent model with a strictly bifurcating topology separating Indian and Holarctic Asian lineages and latter southwestern Asia from central Asia populations, (2) an introgression model with the same strictly bifurcating topology followed by an introgression event from the Indian to the southwestern Asian population (3) an hybridization model, where southwestern Asian population was originated by hybridization between Indian and central Asian populations. Input files including template and parameter estimation files for the three models, are available at **Figure S27.** A summary of all defined parameters and their search ranges are given in **Table S4.**

We performed genotype calling using BCFtools specifically keeping monomorphic sites (Danecek et al. 2021). We filtered our VCF containing all variant and monomorphic sites by the following parameters, keeping only those loci with >10x coverage, and > 50% missing data and removing sites SNPs violating Hardy-Weinberg equilibrium for heterozygous excess (p < 0.01). After filtering, we retained 2,771,059 SNPs and 456,437 monomorphic sites, which we used to generate the joint 3D-SFS using the (folded) minor allele frequency spectrum. We used the canFam 3.1 domestic dog assembly as the reference, resulting in the reference being more closely related to Holarctic wolves than to Indian wolves. Thus, we could not determine the ancestral state of each allele and therefore used the minor allele frequency spectrum (folded SFS). The mutation rate was set to 4.5 x 10^-9^ based on de novo mutations in a pedigree of wolves (Koch et al. 2019). To generate the 3D-SFS while ensuring no missing data and accounting for local linkage disequilibrium (LD) patterns, we down-sampled individuals. Given an estimated recombination rate of 1.34cM/Mb using the recombination map of the dog genome (Auton et al. 2013), we partitioned the genome into independent and unlinked blocks of 2Mb, ensuring that sites within blocks were linked, while those between blocks were unlinked. Within each block obtained, we retained only SNPs without missing data by sampling three, six, and nine individuals from the Indian, southwest, and central populations, respectively, ensuring that each sampled individual had no missing data for that block. FASTSIMCOAL2 estimates a composite likelihood, assuming independence between sites (i.e., it does not account for linkage). Consequently, although the maximized likelihood converges to the correct parameter values, the estimated likelihood approximates the true value when derived from a set of unlinked loci (Excoffier et al., 2013). We utilized the parameter estimates inferred from all retained SNPs to recalculate the likelihoods for each model using only one SNP per block. This recalculation was based on a 3D-SFS derived from a set of potentially independent SNPs, with one SNP selected per block. We used this newly computed likelihood to apply the Akaike Information Criterion (AIC) to compare the three models and computed the relative likelihood of models. For this we calculated delta AIC according to Excoffier et al (2013), which represents the difference between the AIC value of each model and the minimum AIC value among all models. Further we thoroughly examined the parameter values that optimized the likelihood for each model. Specifically, we compared divergence times between Holarctic and Indian lineages, the effective population sizes of each population, and the admixture proportions estimated under each model. This analysis aimed to determine whether the parameter estimates that optimized each model were consistent with the most likely model as indicated by the AIC. In addition, to evaluate whether the best-selected model could accurately reproduce the observed data, we conducted a visual examination comparing the expected and observed marginal one-dimensional SFS, as well as various two-dimensional SFS pairs.

### Long-term demographic history with PSMC

To infer the historical demography of wolves in Eurasia, we used the Pairwise Sequential Markovian Coalescent (PSMC; Li and Durbin 2011). PSMC uses a coalescent approach to estimate the history of change in effective population sizes over time. We only included the autosomal sequences of each gray wolf individual. We converted each bam file to a fasta-like consensus sequence by first using the mpileup command with SAMtools and subsequently using BCFtools view –c to call variants and vcfultils.pl vcf2fq to convert the vcf file to fastq format (Danecek et al. 2021). We excluded any reads that were less than 20 for minimum mapping quality and minimum base quality (-q 20 -Q 20) and excluded reads with excessive mismatches (-C 50). We also removed sites with more than double or less than a third of the average depth of coverage for each sample. We tested different combinations of parameters to infer the PSMC, which were “`psmc -N25 -t15 -r5 -p 4+25*2+4+6`”, “`psmc -N25 -t15 -r5 -p 2+2+25*2+4+6`” and “`psmc -N25 -t15 -r5 -p 1+1+1+1+25*2+4+6`” following previous studies on gray wolves and recent recommendations for inferring PSMC trajectories (Hilgers et al. 2024). Some samples showed a false peak when using the parameter “`psmc -N25 -t15 -r5 -p 4+25*2+4+6`”, therefore, we used the parameter “`psmc -N25 -t15 -r5 -p 1+1+1+1+25*2+4+6`” for our final analyses (**Figure S25**).

To account for our selected low coverage genomes (15-20x), we estimated the false negative rate (FNR) by a downsampling high coverage gray wolf genome that were in the same geographic region to the specific depth of the low coverage genome. To determine the best FNR, we visually estimated the best correspondence between the PSMC plots with the high coverage (>20x) regional wolf genome and downsampled genomes with various FNR corrections (**Figure S26**). We then applied the best estimated FNR to the low coverage gray wolf genomes to infer their demographic history. We used a mutation rate of 4.5 x 10^-9^ (Koch et al. 2019) and a generation time of 4.4 years (Mech and Barber-Meyer 2017).

### Genetic Diversity and Runs of Homozygosity

We calculated the genome-wide heterozygosity for each wolf individual above 5x using ANGSD (Korneliussen et al. 2014). We first estimated the folded site allele frequency with the realSFS program in ANGSD (-doSaf) using genotype likelihoods with the SAMtools model (-GL 1) and using the Andean fox genome to determine ancestral state. We filtered each individual bam file to keep only autosomal chromosomes, only properly paired reads (-only_proper_paired 1), excluded reads with excess of mismatches (-C 50), excluded reads with a mapping quality (-minMapQ 20), excluded reads with a depth less than 4 (-setMinDepth 4), and kept bases with a sequence quality above 20 (-minQ 20). To calculate the genome-wide heterozygosity, we divided the number of heterozygous sites by the total number of sites for each sample.

We inferred runs of homozygosity (ROH) using BCFtools/RoH (Narasimhan et al. 2016). Along with the GATK filtering criteria, we included only biallelic SNPs that had all individuals present per site (--max-missing 1, --min-alleles 2, --max-alleles 2). We also included SNPs that had a depth of more than one third and less than double of the average depth of coverage for each sample, where we chose samples that were >15x. For the BCFtools/RoH analysis, we fixed the alternative allele frequency to 0.4 (-AFdflt 0.4) and used the dog recombination map from Auton et al. 201x to account for recombination hotspots (--genetic-map). We kept ROHs that had a quality score of at least 80 and excluded ROHs that were below 100 kb in length in our analysis.

To gain insight into the timing that inbreeding occurred, we calculated the expected generations since inbreeding using the length of the ROHs, generation time of 4.4 years (Mech and Barber-Meyer 2017), and an average recombination rate of 1.34 cM/Mb calculated as the average recombination rate across the autosomes from the canfam3.1 genome assembly (Auton et al. 2013). Specifically, we used the formula: g=100(2rLROH) where g=generation interval, r=recombination rate (cM/Mb), and L=Length of ROH (Mb) (Thompson et al. 2013). For example, a ROH length of 5Mb will have an expected time of coalescence of 7.46 generations ago, corresponding to inbreeding occurring around 32 years ago using a wolf generation time of 4.4 years. Likewise, we estimate that a ROH length of <0.16 Mb corresponds to inbreeding occurring >1,026 years ago.

### Inbreeding coefficient

We inferred the per-individual inbreeding coefficient (Find) using genotype likelihoods, where the inbreeding coefficient is the probability of identity by descent. We used ANGSD to infer genotype likelihoods and allele frequencies for samples above 4x coverage with the following parameters: -GL 2 -checkBamHeaders 0 -trim 0 -C 50 -baq 1 -skipTriallelic 1 -minMapQ 20 -minQ 20 -minInd 88 -uniqueOnly 1 -remove-bads 1 - noTrans 1 -doGlf 3 -doMajorMinor 1 -doMaf 1 -minMaf 0.0001 -SNP_pval 1e-6. We then used NgsRelate v2 with option -F1 to infer the inbreeding coefficients for each Eurasian wolf individual (Hanghoj et al. 2019).

### Genetic Load

We selected 14 individuals with > 20x sequencing coverage from India (n=2), the Tibetan plateau (n=2), southwestern Asia (n=4), Central Asia (n=4), and the Mexican wolf (n=1). We selected these wolves to assess the genetic load of Indian and Tibetan wolves, in which we compared Indian and Tibetan wolf genetic load with two adjacent, genetically diverse populations, and to a Mexican wolf individual with previously documented very low genetic diversity. We used SnpEff to annotate a VCF, including only biallelic non-indel sites with no missing data, minimum depth of 10, maximum depth of 60, a minimum allele count of 2 and a minimum sequence quality (--minQ) of 30.

To polarize our dataset, we followed the method by Smeds et al. (2022) to map two outgroups to the dog reference genome at only sites with a minimum depth of 4 at both outgroup and included nonvariant sites. We used Dhole (*Cuon alpinus*) and Andean fox (*Lycalopex culpaeus*) as our outgroups. We then performed pseudo-haploidization of the outgroup genomes and kept only sites in which the allele agreed in both outgroups (Smeds et al. 2022). We used this to add the ancestral allele information to our annotated VCF, and assumed that the derived allele was the deleterious allele (Smeds et al. 2022).

We counted the number of derived alleles and whether they were in a homozygous or heterozygous state for variants predicted by SnpEff to have high, moderate, and low impact on protein function. High impact is predicted to cause loss of function or protein truncation (stop gain, frameshift mutation); moderate impact is predicted to change protein effectiveness (missense variant), and low impact is assumed harmless, such as a synonymous variant. We also calculated the following metrics: total load (the total number of derived alleles classified as high or moderate impact), realized load (homozygous state of derived alleles in high or moderate impact), and masked load (heterozygous state of derived alleles in high or moderate impact).

To investigate evidence of differential loss of derived variants for different types of mutations, we assessed the relative amounts of mutational categories given by SnpEff. These mutational categories are: Loss of Function (LOF) mutations that imply a high predicted impact on protein structure, missense mutations (t moderate impact), synonymous mutations (low impact), and “modifier” mutations found in non-coding regions (no impact). We calculated the count of derived alleles and the number of heterozygous and homozygous derived allele calls in each of the four mutational categories for each wolf individual. To account for differences in overall derived allele counts among individuals, we calculated the average derived allele count per 100,000 derived neutral alleles for total mutational load, LOF, missense, and synonymous mutations. We considered the modifier category as neutral derived alleles. Specifically, we calculated mutational load as: (total derived alleles in LOF and missense / total derived alleles in modifier category) multiplied by 100,000.

## Acknowledgements

L.M.H thanks the National Science Foundation Postdoctoral Research Fellowship (award number 2208950) for funding and support. The Norwegian Environment Agency (project 18088069) provided funding and support for sequencing efforts on newly sequenced wolf genomes. Ç.H.Ș. thanks Fondation Segré and the Sigrid Rausing Trust for providing the majority of the funding for this project, H. Batubay Özkan and Barbara Watkins for their support of the Biodiversity and Conservation Ecology Lab at the University of Utah, and Bilge Bahar, Seha İşmen, Ömer Koç, Ömer Külahçıoğlu, Burak Över, Emin Özgür, Suna Reyent and Ceren Sağlamer for supporting this project. Türkiye’s Department of Nature Conservation and National Parks and the Ministry of Agriculture and Forestry granted the permit for Türkiye (No. 72784983-488.04-114100). We thank the Museum of the Institute of Plant and Animal Ecology UB RAS access to their collections.

## Author Contributions

L.M.H. conceived and designed the project with guidance from S.G., M.H.S.S., and B.N.S. C.S.O., M.H.S.S., N.F.G.M. conducted laboratory work, H.F., G.S., B.H., F.H., S.P.E.H., C.K., Ç.H.Ş., P.K., H.S., M.H.S.S., M.T.P.G. provided logistics, field work, wolf samples, and sequencing efforts, L.M.H. led data analysis with assistance from B.R.P., A.N., X.S., N.F.G.M, L.M.H. wrote the manuscript with input from all co-authors.

## Data Availability

All raw reads have been submitted to the NCBI Project ID XXXX, with accession numbers XXXX-XXXX. The bioinformatic scripts and results from the phylogenetic analyses, D statistics, genetic load, and PSMC can be found on the github: https://github.com/hennelly/Asia_wide_wolf_genomics.

