## Supplemental Figures for "Complex genomic ancestry in southern regions and drivers of continental-level genetic diversity in the wolves of Asia"

**Figure S1.** The estimated distribution of the three extant divergent wolf lineages – the Indian, Tibetan, and Holarctic lineage
**Figure S2.** Individual admixture proportions using K=2 to K=8 populations for 115 individuals consisting of various canid species, wolves, and dogs
**Figure S3.** D statistic values to assess derived allele sharing between each wolf individual in Eurasia (X) with the Tibetan wolf with the topology: (Graywolf$_{Norway}$, X), Tibetan wolf–TI32), Andeanfox).
**Figure S4.** Derived allele sharing between each wolf individual in Eurasia (circles) with the Tibetan wolf with the topology: (Graywolf$_{Norway}$, X), Tibetan wolf–TI32), Andeanfox).
**Figure S5.** Phylogeny of the autosomes inferred with ASTRAL 5.7.8, using 1000 genomic regions at a 20kb length (Zhang et al. 2018).
**Figure S6**. Fully labeled phylogenetic tree of the X chromosome using 514,048 SNPs found in only the low recombination regions (<0.2cM/Mb).
**Figure S7**. Fully labeled phylogenetic tree of the X chromosome using 514,048 SNPs found in only the low recombination regions (<0.2cM/Mb) where we partitioned this set of SNPs into 10kb genomic regions.
**Figure S8**. Fully labeled phylogenetic tree using only low recombination regions (<0.2cM/Mb) across the autosomes, where we inferred the phylogeny by partitioned 6.06 million SNPs into 10kb genomic regions.
**Figure S9.** Phylogeny of the X chromosome inferred with ASTRAL 5.7.8, using 500 genomic regions at a 20kb length (Zhang et al. 2018).
**Figure S10.** (A) Average topology weight across the X chromosome for Eastern Asian wolves and wolves from the Central Asian mountains within three categories: low recombination regions (<0.2cM/Mb), medium (0.2-2cM/Mb), and high (>2cM/Mb).
**Figure S11.** D statistic values to assess derived allele sharing between each wolf individual in Eurasia (X) with African wolves with the topology: (Graywolf$_{Norway}$, X), African wolf), Andeanfox).
**Figure S12.** Derived allele sharing between each wolf individual by population with the Indian wolf and the African wolf.
**Figure S13.** Derived allele sharing between each wolf individual in Eurasia (circles) with the Indian wolf with the topology: (Graywolf$_{Norway}$, X), Indian wolf–BH123), Andeanfox).
**Figure S14.** D statistic values to assess derived allele sharing between each wolf individual in Eurasia (X) with the Indian wolf with the topology: (Graywolf$_{Norway}$, X), Indian wolf BH123), Andeanfox).
**Figure S15.** Three alternative scenarios for the origin of the Southwest Asian population tested in fastsimcoal2 (Excoffier et al. 2021).
**Figure S16**. Fully labeled phylogenetic tree of the X chromosome using 514,048 SNPs found in only the low recombination regions (<0.2cM/Mb) with excluding wolves from Pakistan and two wolves from southwest Asia that were clustered with Indian wolves (wolf from Zanjan and Hamadan in Iran).
**Figure S17**. (A,B,C) Maximum likelihood autosomal phylogeny of wild canids inferred with IQ-Tree 1.6.12 using only low recombination (<0.2cM/Mb) regions of the X chromosome, we used

IQ-Tree 1.6.12 where we estimated the best model using ModelFinder and used 1,000 ultra-fast bootstraps to infer each tree.

**Figure S18**. **(A)** Fully labeled maximum likelihood phylogeny using all wild canid individuals excepts dogs and wolves from Pakistan and Ladakh inferred using autosomal genomic regions in which the topology weight of topology 73 was above 0.3 (~99.99$^{th}$ percentile; ~40kb length total).

**Figure S19**. **(A)** A maximum likelihood phylogeny using all wild canid individuals excepts dogs and wolves from Pakistan and Ladakh inferred using autosomal genomic regions in which the topology weight of topology 73 was above 0.2 (~99.5$^{th}$ percentile; ~299kb length total).

**Figure S20**. Estimated inbreeding coefficients ($F_{Ind}$) using genotype likelihoods with NgsRelate for 98 individuals across seven wolf populations in Eurasia.

**Figure S21.** Timing of inbreeding estimated from the length of ROH blocks across the autosomes of 42 wolves.

**Figure S22.** Number of heterozygous and homozygous counts for the derived allele for High impact, Medium impact, and Low impact categories. Homozygous counts contain two derived alleles and heterozygous derived counts contain one derived allele.

**Figure S23.** Total genetic load (the total number of derived alleles; homozygous derived alleles counted twice, heterozygous derived alleles once), realized load (homozygous state of derived alleles), and masked load (heterozygous state of derived alleles) for each impact category (High, Moderate, Low) for five selected wolf populations of wolves.

**Figure S24**. Cross validation results by using leave-one-out cross validation to select the optimal value of lambda, the smoothing parameter.

**Figure S25.** PSMC plots for three wolf genomes while varying the atomic time intervals.

**Figure S26**. Down-sampling of genomes to select the false negative rates (FNR) for low-depth corrections of pairwise sequentially Markovian coalescent (PSMC) demographic trajectories.

**Figure S27.** Est and Tpl input files for fastsimcoal2 for each of the three models we tested.

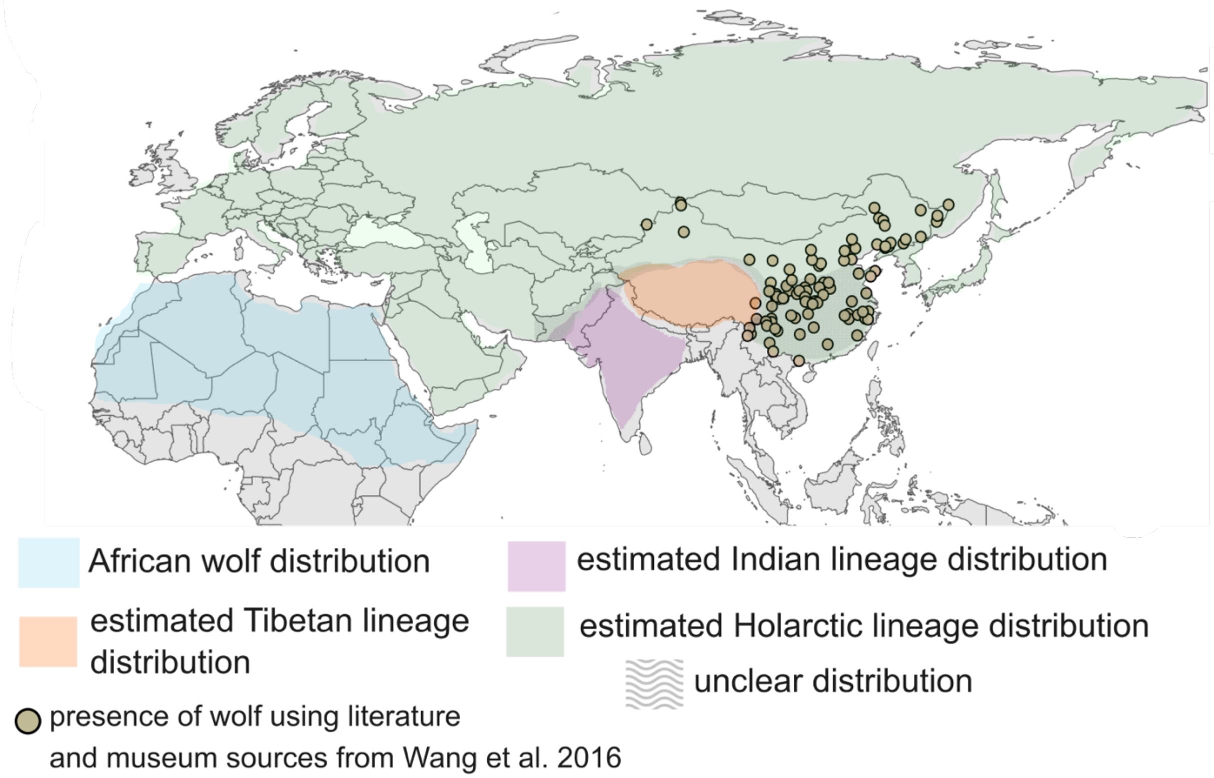


**Figure S1.** The estimated distribution of the three extant divergent wolf lineages – the Indian, Tibetan, and Holarctic lineage. The estimated distribution is based on previous work on geographic distribution of mitochondrial haplotypes, where three major mitochondrial lineages are present in extant wolves (Ersmark et al. 2016, Werhahn et al. 2023, Hennelly et al. 2023, Werhahn et al. 2020, Hoffmann and Atickem 2019). Previous studies found concordance between mitochondrial haplotype and nuclear genomic ancestry for the three wolf lineage (Werhahn et al. 2020, Wang et al. 2020, Hennelly et al. 2021, Hennelly et al. 2023). Wavy lines indicate unclear distribution due to lack of data. Circles indicate locations of wolf presence based on literature sources and museum specimens within Table 1 and Table 2 of Wang et al. 2016. Records of wolves are found throughout China, including southern regions bordering Vietnam. Wavy lines are indicated in eastern and southern China due to lack of samples that have been sequenced from these regions and lack of resolution into their lineage assignment (Wang et al. 2019).

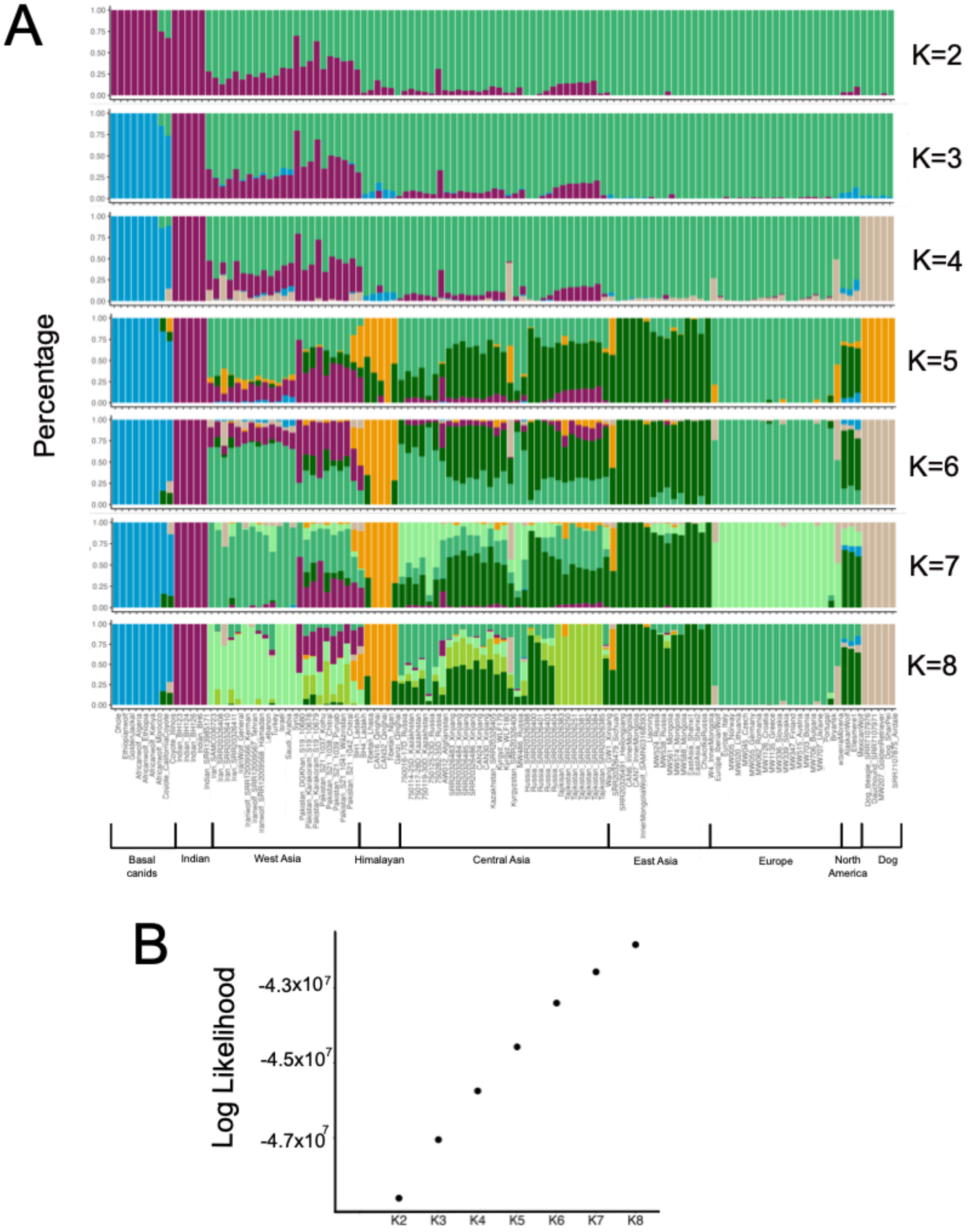

**Figure S2.** Individual admixture proportions using K=2 to K=8 populations for 115 individuals consisting of various canid species, wolves, and dogs. We used 578,343 SNPs inferred using genotype likelihoods with ANGSD, which were used to estimate the individual admixture proportions through NGSAdmix (Skotte et al. 2013, Korneliussen et al. 2014). **(A)** Each bar represents an individual and colors within each bar represent the estimated ancestry belonging to a specific ancestry. **(B)** Log likelihoods of each run in NGSAdmix.

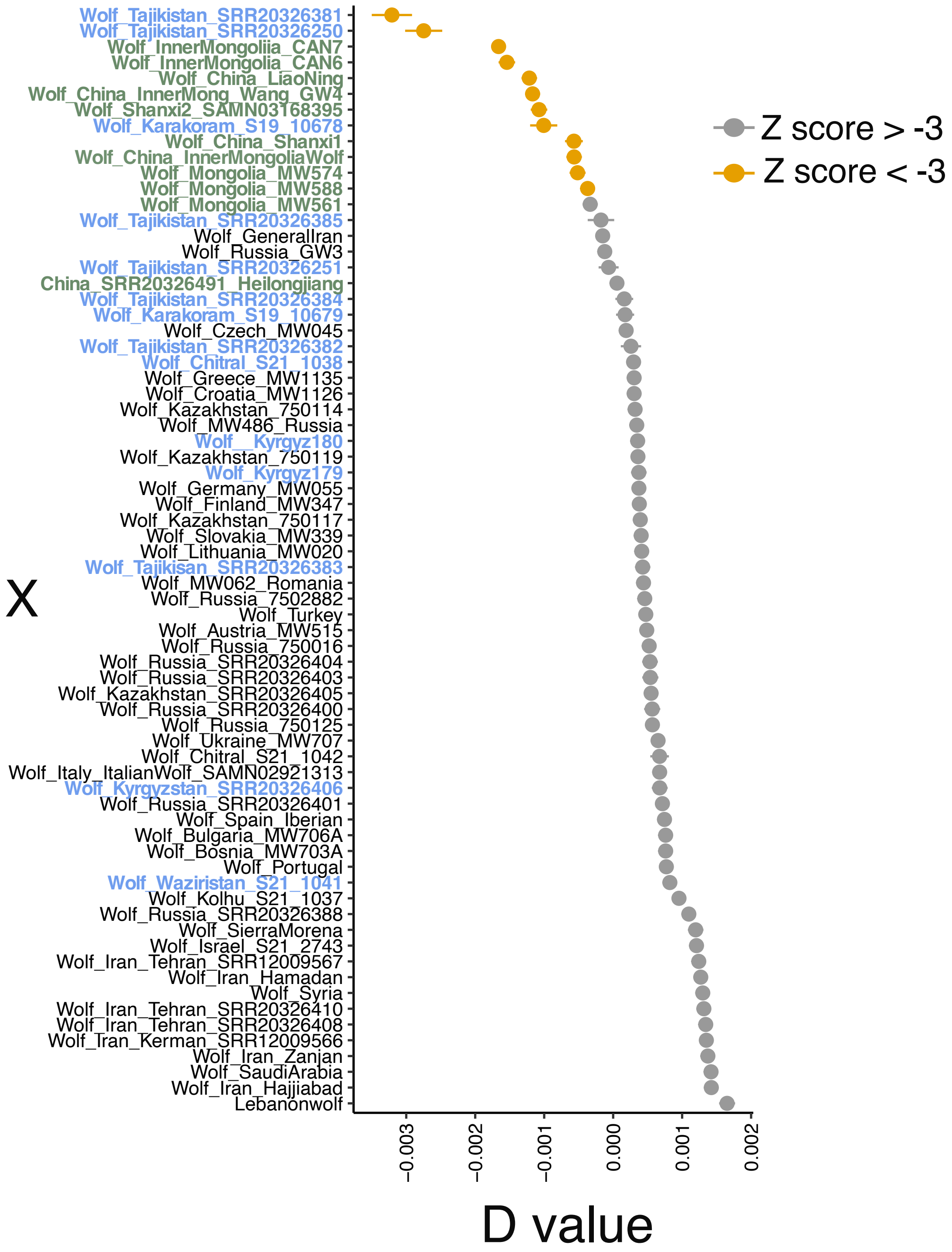


**Figure S3.** D statistic values to assess derived allele sharing between each wolf individual in Eurasia (X) with the Tibetan wolf with the topology: (Graywolf$_{Norway}$, X), Tibetan wolf–TI32), Andeanfox). A negative D-value indicates an excess derived allele sharing with the Tibetan wolf, while a positive D value can indicate allele sharing between X and the Andean fox, or X and the Gray wolf from Norway. Yellow color indicates D values that show statistically significant derived allele sharing with the Tibetan wolf (Z-score < -3). Green and blue labeled wolf samples indicate wolves that from the Eastern Asia and Central Asian mountains, respectively.

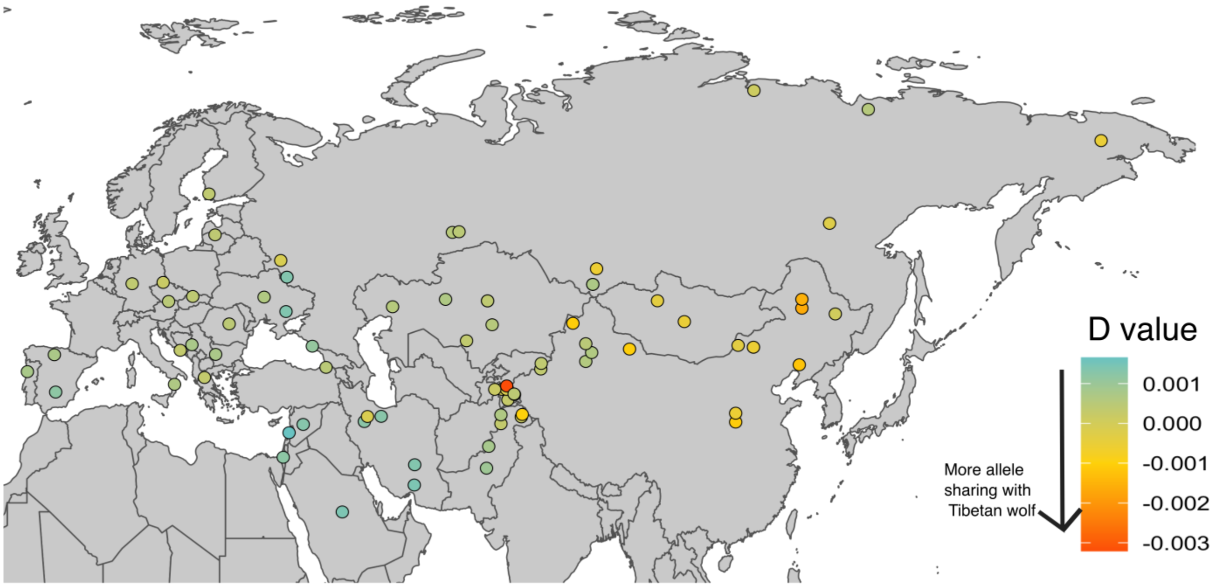


**Figure S4.** Derived allele sharing between each wolf individual in Eurasia (circles) with the Tibetan wolf with the topology: (Graywolf$_{Norway}$, X), Tibetan wolf–TI32), Andeanfox). A negative D-value indicates an excess derived allele sharing with the Tibetan wolf, while a positive D value can indicate allele sharing between X and the Andean fox, or X and the Gray wolf from Norway.

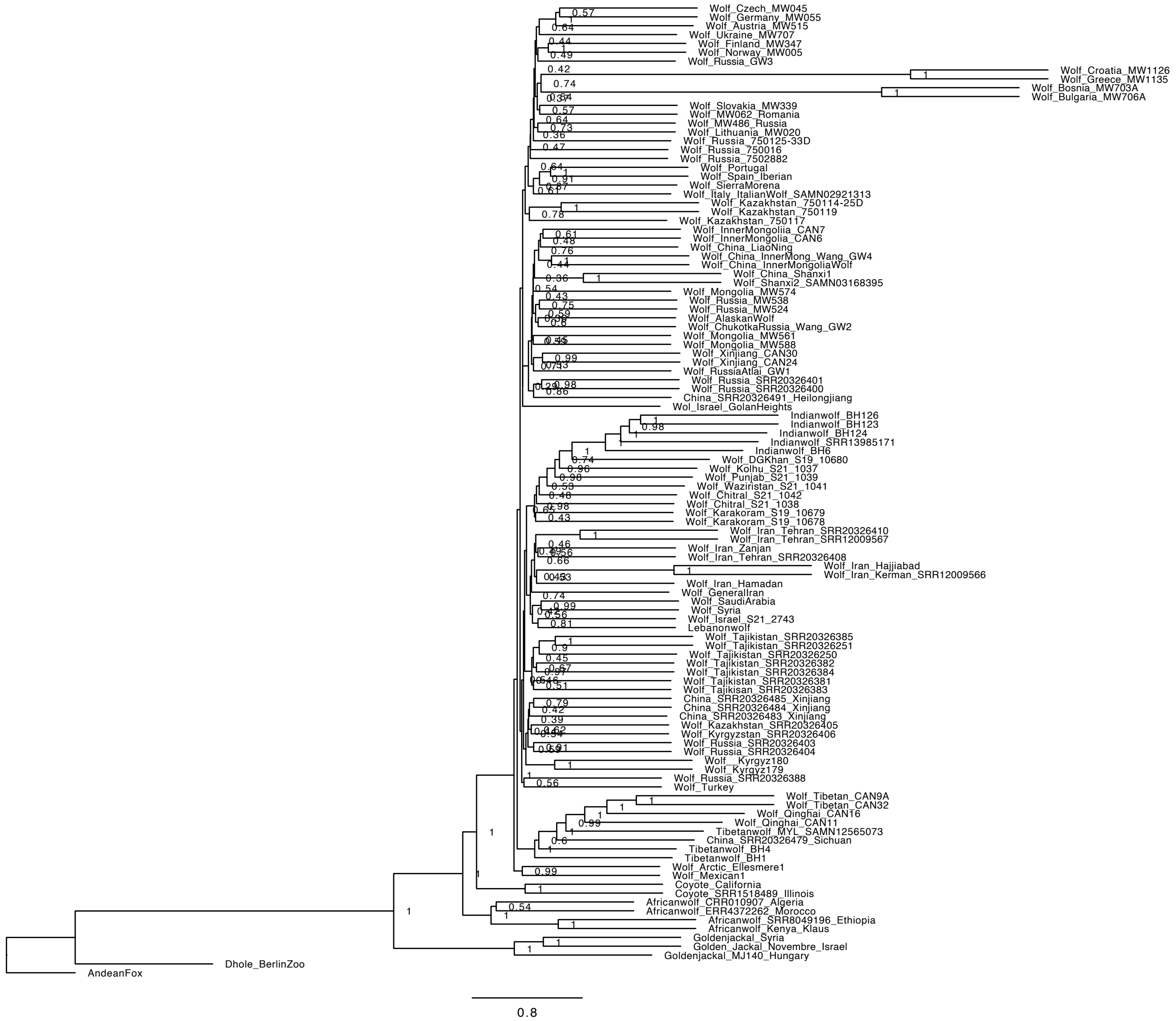


**Figure S5.** Phylogeny of the autosomes inferred with ASTRAL 5.7.8, using 1000 genomic regions at a 20kb length (Zhang et al. 2018). We converted each 20kb VCF file to a phylip file using the vcf2phy.py script. We inferred a maximum-likelihood phylogenetic tree for each 20kb region using IQ-Tree 1.6.12 where we estimated the best model using ModelFinder and used 1,000 ultra-fast bootstraps to infer each tree (Kalyaanamoorthy et al. 2017, Nguyen et al. 2014). Each node is labeled with the local posterior probability (LPP), which are computed based on gene tree quartet frequencies where a higher LPP has lower discordance.

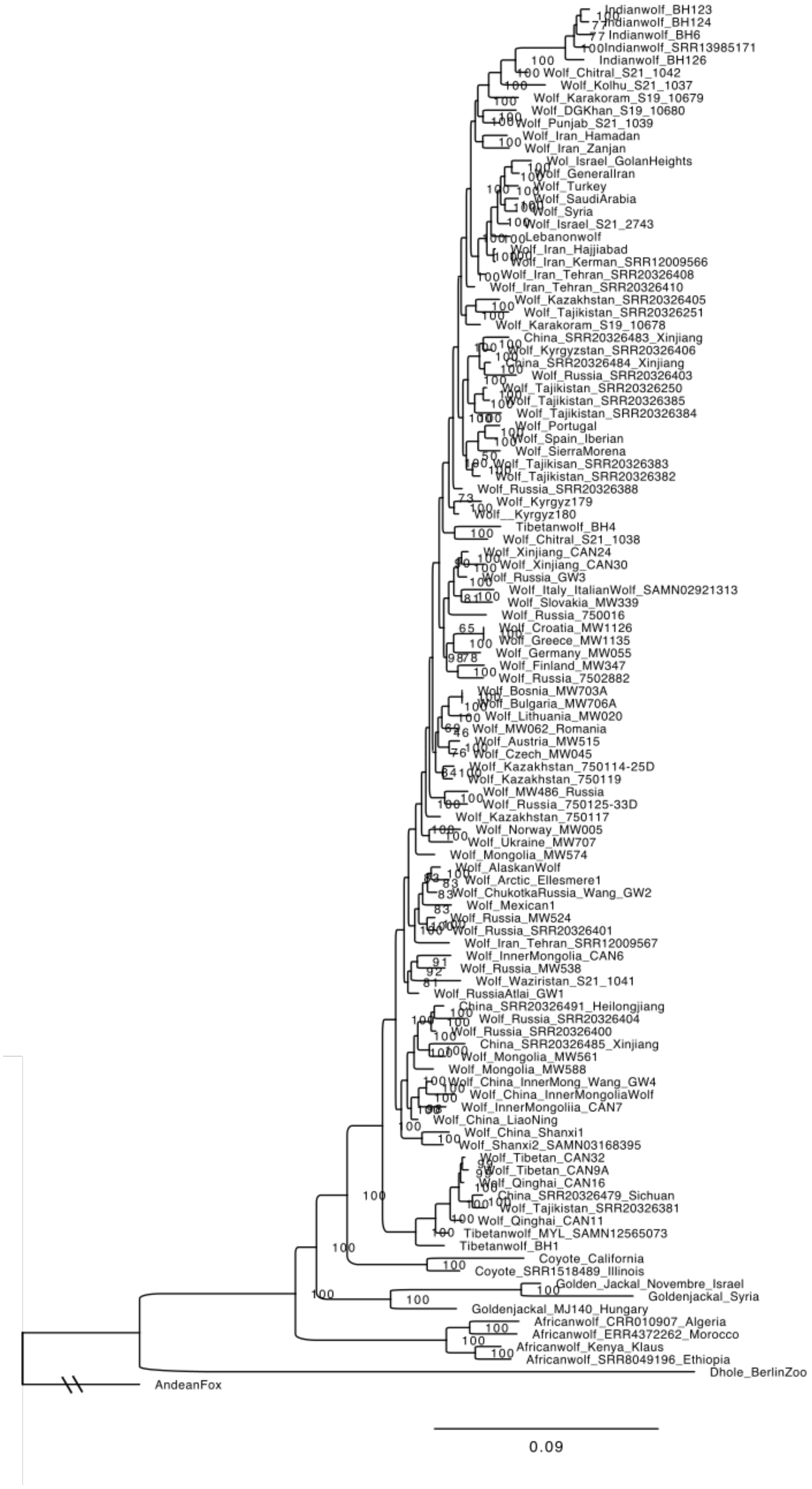
Indianwolf_BH123
Indianwolf_BH124
Indianwolf_BH6
Indianwolf_SRR13985171
Indianwolf_BH126
Wolf_Chitral_S21_1042
Wolf_Kolhu_S21_1037
Wolf_Karakoram_S19_10679
Wolf_DGKhan_S19_10680
Wolf_Punjab_S21_1039
Wolf_Iran_Hamadan
Wolf_Iran_Zanjan
Wol_Israel_GolanHeights
Wolf_GeneralIran
Wolf_Turkey
Wolf_SaudiArabia
Wolf_Syria
Wolf_Israel_S21_2743
Lebanonwolf
Wolf_Iran_Hajjiabad
Wolf_Iran_Kerman_SRR12009566
Wolf_Iran_Tehran_SRR20326408
Wolf_Iran_Tehran_SRR20326410
Wolf_Kazakhstan_SRR20326405
Wolf_Tajikistan_SRR20326251
Wolf_Karakoram_S19_10678
China_SRR20326483_Xinjiang
Wolf_Kyrgyzstan_SRR20326406
China_SRR20326484_Xinjiang
Wolf_Russia_SRR20326403
Wolf_Tajikistan_SRR20326250
Wolf_Tajikistan_SRR20326385
Wolf_Tajikistan_SRR20326384
Wolf_Portugal
Wolf_Spain_Iberian
Wolf_SierraMorena
Wolf_Tajikisan_SRR20326383
Wolf_Tajikistan_SRR20326382
Wolf_Russia_SRR20326388
Wolf_Kyrgyz179
Wolf__Kyrgyz180
Tibetanwolf_BH4
Wolf_Chitral_S21_1038
Wolf_Xinjiang_CAN24
Wolf_Xinjiang_CAN30
Wolf_Russia_GW3
Wolf_Italy_ItalianWolf_SAMN02921313
Wolf_Slovakia_MW339
Wolf_Russia_750016
Wolf_Croatia_MW1126
Wolf_Greece_MW1135
Wolf_Germany_MW055
Wolf_Finland_MW347
Wolf_Russia_7502882
Wolf_Bosnia_MW703A
Wolf_Bulgaria_MW706A
Wolf_Lithuania_MW020
Wolf_MW062_Romania
Wolf_Austria_MW515
Wolf_Czech_MW045
Wolf_Kazakhstan_750114-25D
Wolf_Kazakhstan_750119
Wolf_MW486_Russia
Wolf_Russia_750125-33D
Wolf_Kazakhstan_750117
Wolf_Norway_MW005
Wolf_Ukraine_MW707
Wolf_Mongolia_MW574
Wolf_AlaskanWolf
Wolf_Arctic_Ellesmere1
Wolf_ChukotkaRussia_Wang_GW2
Wolf_Mexican1
Wolf_Russia_MW524
Wolf_Russia_SRR20326401
Wolf_Iran_Tehran_SRR12009567
Wolf_InnerMongolia_CAN6
Wolf_Russia_MW538
Wolf_Waziristan_S21_1041
Wolf_RussiaAtlai_GW1
China_SRR20326491_Heilongjiang
Wolf_Russia_SRR20326404
Wolf_Russia_SRR20326400
China_SRR20326485_Xinjiang
Wolf_Mongolia_MW561
Wolf_Mongolia_MW588
Wolf_China_InnerMong_Wang_GW4
Wolf_China_InnerMongoliaWolf
Wolf_InnerMongoliia_CAN7
Wolf_China_LiaoNing
Wolf_China_Shanxi1
Wolf_Shanxi2_SAMN03168395
Wolf_Tibetan_CAN32
Wolf_Tibetan_CAN9A
Wolf_Qinghai_CAN16
China_SRR20326479_Sichuan
Wolf_Tajikistan_SRR20326381
Wolf_Qinghai_CAN11
Tibetanwolf_MYL_SAMN12565073
Tibetanwolf_BH1
Coyote_California
Coyote_SRR1518489_Illinois
Golden_Jackal_Novembre_Israel
Goldenjackal_Syria
Goldenjackal_MJ140_Hungary
Africanwolf_CRR010907_Algeria
Africanwolf_ERR4372262_Morocco
Africanwolf_Kenya_Klaus
Africanwolf_SRR8049196_Ethiopia
Dhole_BerlinZoo
AndeanFox
0.09

**Figure S6**. Fully labeled phylogenetic tree of the X chromosome using 514,048 SNPs found in only the low recombination regions (<0.2cM/Mb). We ran IQ-Tree 1.6.12 with estimating the best model using ModelFinder, which was TVM+F+R4, and used 1,000 ultra-fast bootstraps to infer each tree (Kalyaanamoorthy et al. 2017, Nguyen et al. 2014). The ultra-fast bootstrap support values are shown, where a 95% support corresponds to a probability of 95% the clade is correct.

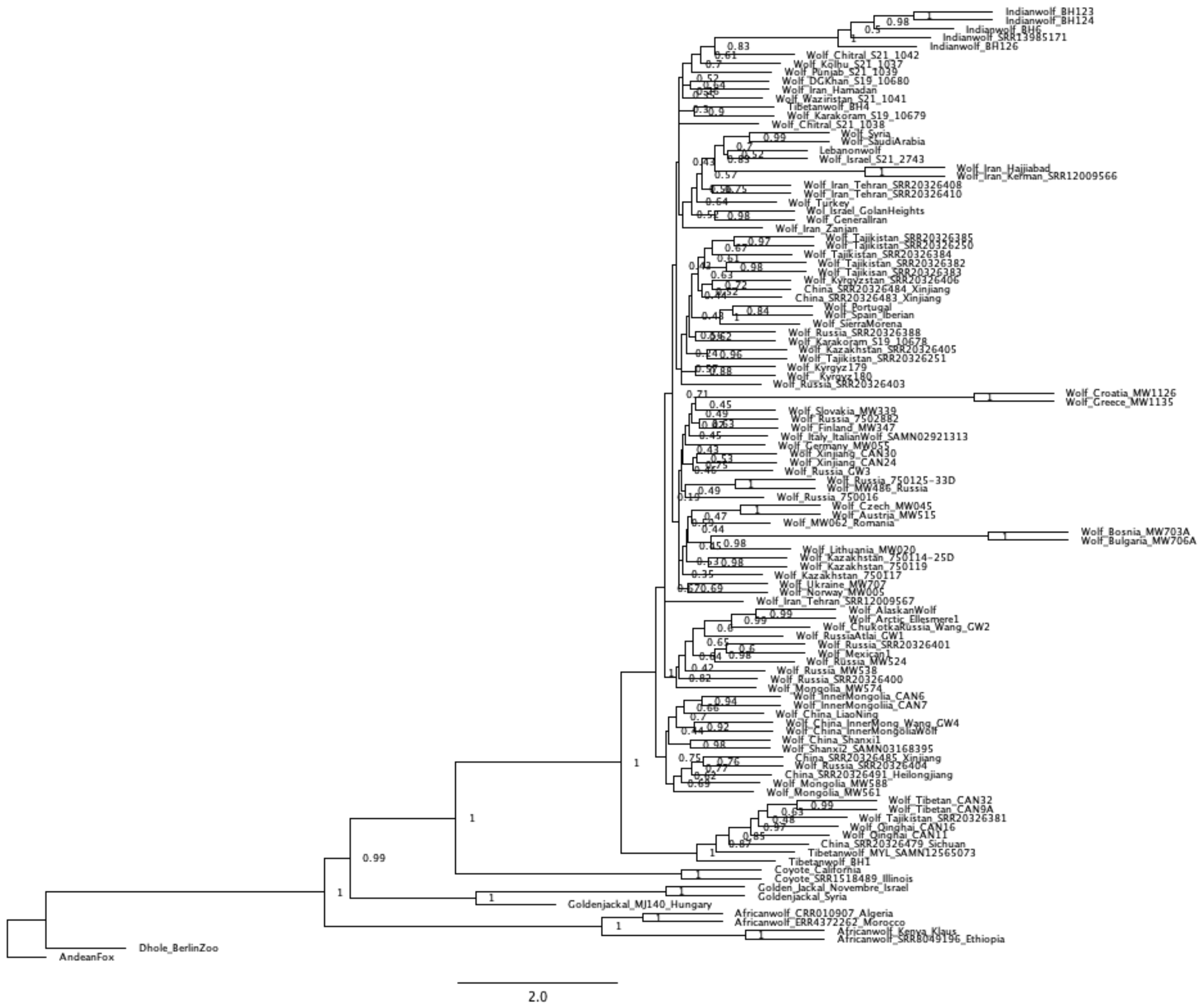


**Figure S7**. Fully labeled phylogenetic tree of the X chromosome using 514,048 SNPs found in only the low recombination regions (<0.2cM/Mb) where we partitioned this set of SNPs into 10kb genomic regions. We ran ASTRAL 5.7.8 using 50 genomic regions that were 10kb in length. Each node is labeled with the local posterior probability (LPP), which are computed based on gene tree quartet frequencies where a higher LPP has lower discordance.

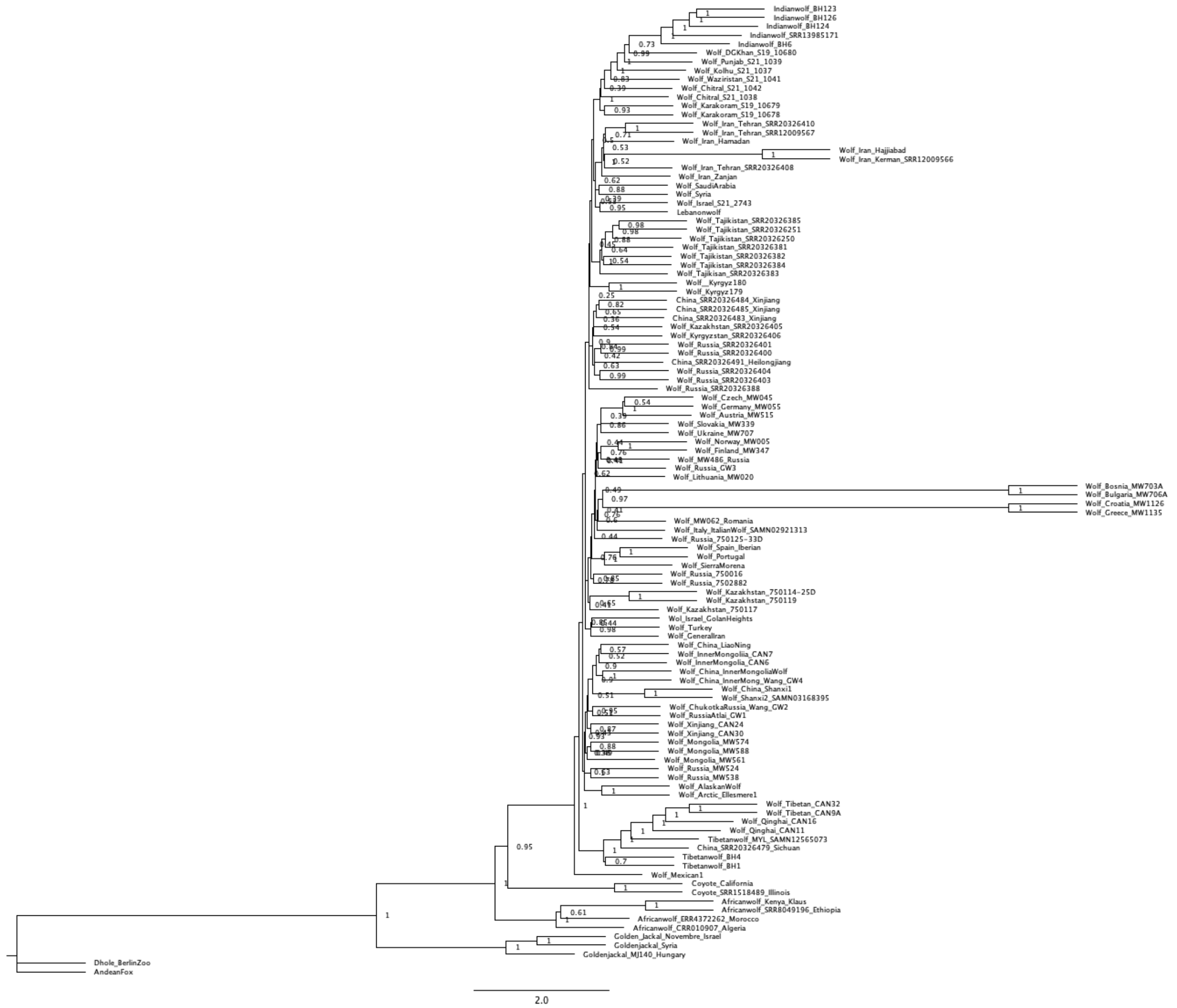


**Figure S8**. Fully labeled phylogenetic tree using only low recombination regions (<0.2cM/Mb) across the autosomes, where we inferred the phylogeny by partitioned 6.06 million SNPs into 10kb genomic regions. We ran ASTRAL 5.7.8 on 606 genomic regions that were 10kb in length. Each node is labeled with the local posterior probability (LPP), which are computed based on gene tree quartet frequencies where a higher LPP has lower discordance.

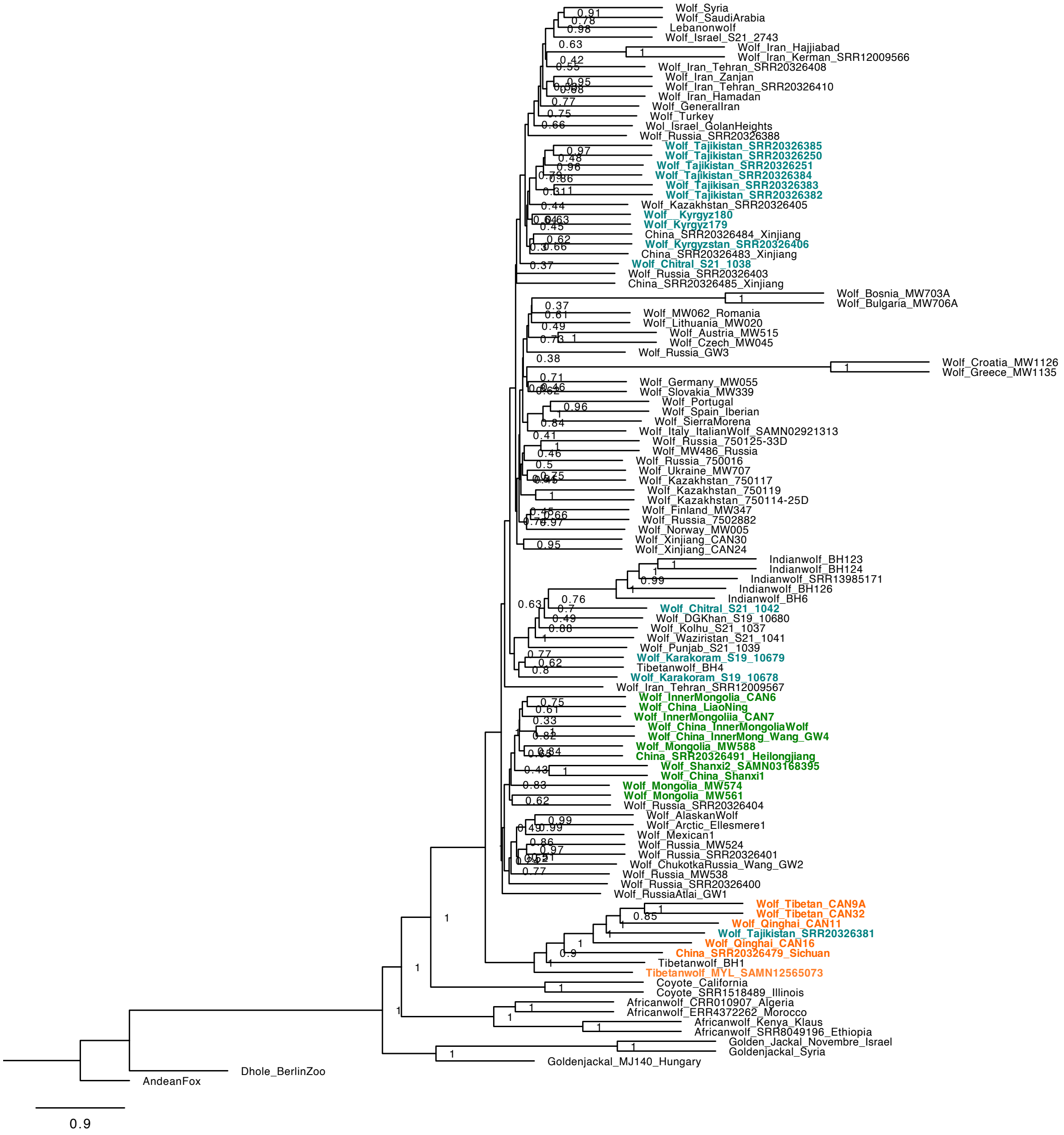

**Figure S9.** Phylogeny of the X chromosome inferred with ASTRAL 5.7.8, using 500 genomic regions at a 20kb length (Zhang et al. 2018). We converted each 20kb VCF file to a phylip file using the vcf2phy.py script. We inferred a maximum-likelihood phylogenetic tree for each 20kb region using IQ-Tree 1.6.12 where we estimated the best model using ModelFinder and used 1,000 ultra-fast bootstraps to infer each tree (Kalyaanamoorthy et al. 2017, Nguyen et al. 2014). Each node is labeled with the local posterior probability (LPP), which are computed based on gene tree quartet frequencies where a higher LPP has lower discordance.

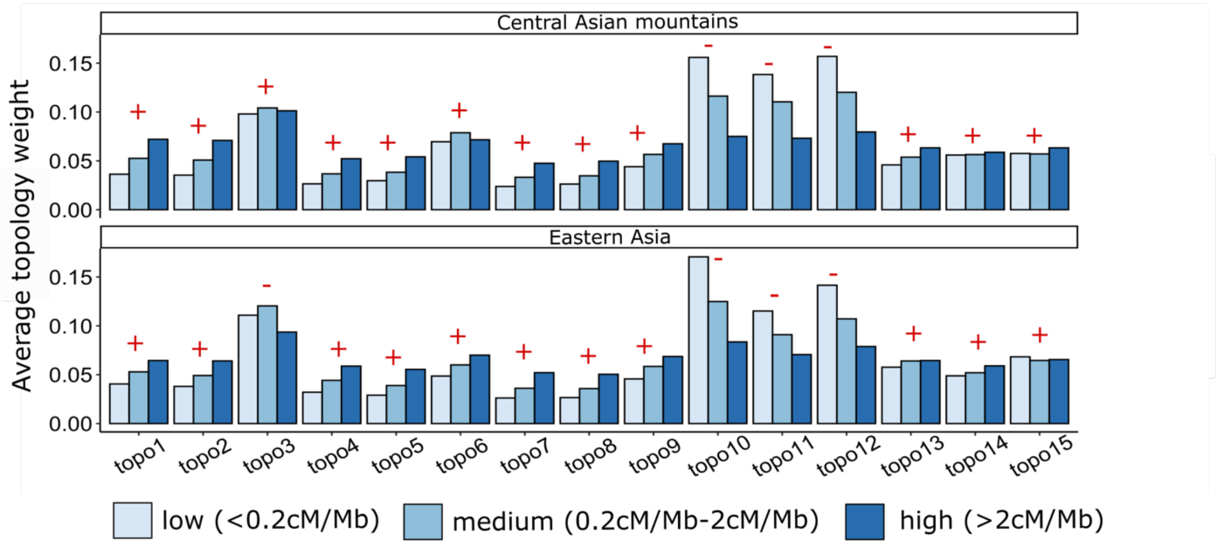


**Figure S10.** (A) Average topology weight across the X chromosome for Eastern Asian wolves and wolves from the Central Asian mountains within three categories: low recombination regions (<0.2cM/Mb), medium (0.2-2cM/Mb), and high (>2cM/Mb). Twisst was ran by (B) Recombination rate variation vs. topology weights of topo10, topo11, topo12 across the X chromosome for Central Asian wolves and Eastern Asian wolves. Signs refer to whether the average within the high recombination region is larger (+) or smaller (-) than the average in the low recombination region.

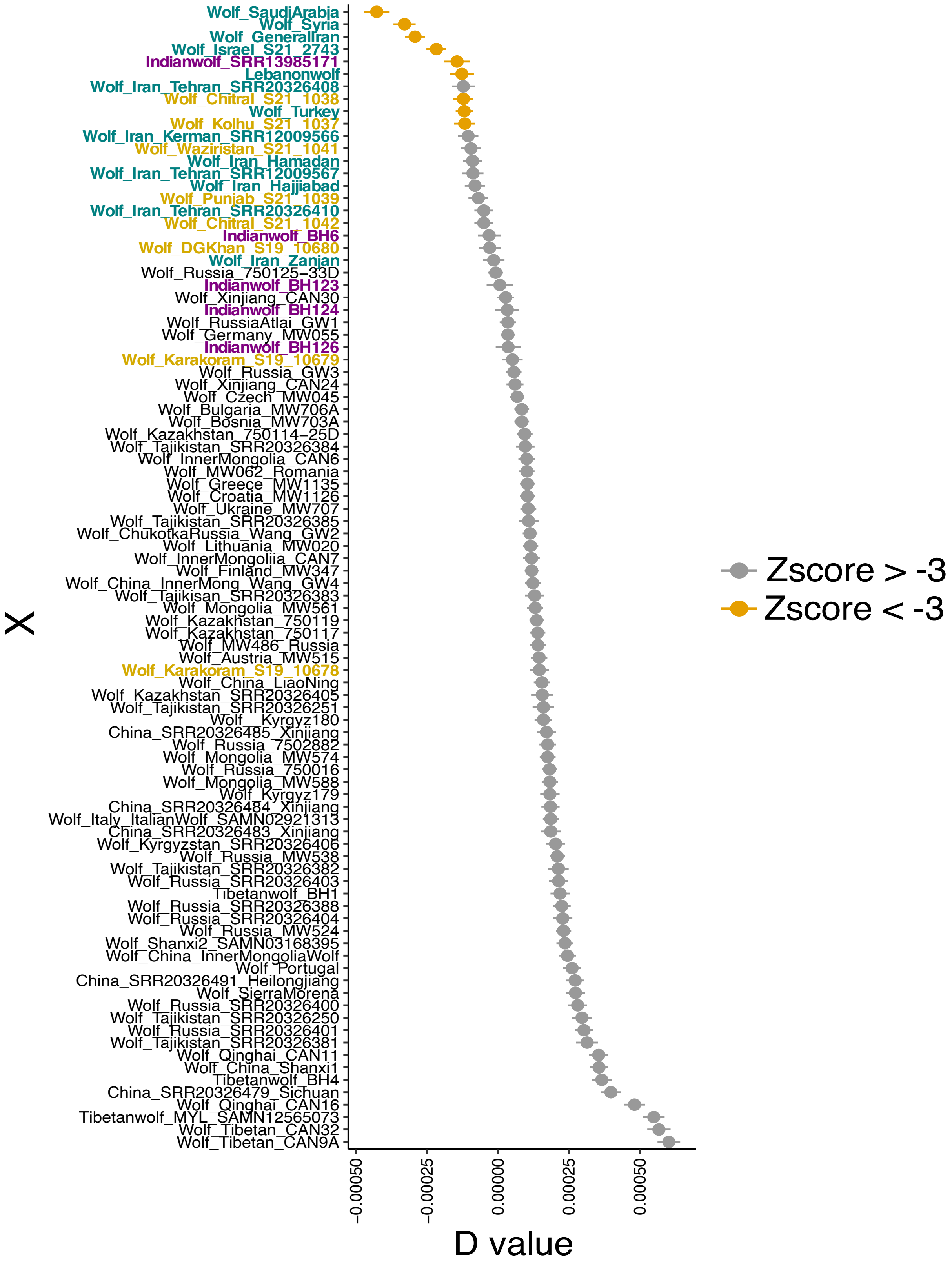
Wolf_SaudiArabia
Wolf_Syria
Wolf_GeneralIran
Wolf_Israel_S21_2743
Indianwolf_SRR13985171
Lebanonwolf
Wolf_Iran_Tehran_SRR20326408
Wolf_Chitral_S21_1038
Wolf_Turkey
Wolf_Kolhu_S21_1037
Wolf_Iran_Kerman_SRR12009566
Wolf_Waziristan_S21_1041
Wolf_Iran_Hamadan
Wolf_Iran_Tehran_SRR12009567
Wolf_Iran_Hajjiabad
Wolf_Punjab_S21_1039
Wolf_Iran_Tehran_SRR20326410
Wolf_Chitral_S21_1042
Indianwolf_BH6
Wolf_DGKhan_S19_10680
Wolf_Iran_Zanjan
Wolf_Russia_750125−33D
Indianwolf_BH123
Wolf_Xinjiang_CAN30
Indianwolf_BH124
Wolf_RussiaAtlai_GW1
Wolf_Germany_MW055
Indianwolf_BH126
Wolf_Karakoram_S19_10679
Wolf_Russia_GW3
Wolf_Xinjiang_CAN24
Wolf_Czech_MW045
Wolf_Bulgaria_MW706A
Wolf_Bosnia_MW703A
Wolf_Kazakhstan_750114−25D
Wolf_Tajikistan_SRR20326384
Wolf_InnerMongolia_CAN6
Wolf_MW062_Romania
Wolf_Greece_MW1135
Wolf_Croatia_MW1126
Wolf_Ukraine_MW707
Wolf_Tajikistan_SRR20326385
Wolf_ChukotkaRussia_Wang_GW2
Wolf_Lithuania_MW020
Wolf_InnerMongoliia_CAN7
Wolf_Finland_MW347
Wolf_China_InnerMong_Wang_GW4
Wolf_Tajikisan_SRR20326383
Wolf_Mongolia_MW561
Wolf_Kazakhstan_750119
Wolf_Kazakhstan_750117
Wolf_MW486_Russia
Wolf_Austria_MW515
Wolf_Karakoram_S19_10678
Wolf_China_LiaoNing
Wolf_Kazakhstan_SRR20326405
Wolf_Tajikistan_SRR20326251
Wolf__Kyrgyz180
China_SRR20326485_Xinjiang
Wolf_Russia_7502882
Wolf_Mongolia_MW574
Wolf_Russia_750016
Wolf_Mongolia_MW588
Wolf_Kyrgyz179
China_SRR20326484_Xinjiang
Wolf_Italy_ItalianWolf_SAMN02921313
China_SRR20326483_Xinjiang
Wolf_Kyrgyzstan_SRR20326406
Wolf_Russia_MW538
Wolf_Tajikistan_SRR20326382
Wolf_Russia_SRR20326403
Tibetanwolf_BH1
Wolf_Russia_SRR20326388
Wolf_Russia_SRR20326404
Wolf_Russia_MW524
Wolf_Shanxi2_SAMN03168395
Wolf_China_InnerMongoliaWolf
Wolf_Portugal
China_SRR20326491_Heilongjiang
Wolf_SierraMorena
Wolf_Russia_SRR20326400
Wolf_Tajikistan_SRR20326250
Wolf_Russia_SRR20326401
Wolf_Tajikistan_SRR20326381
Wolf_Qinghai_CAN11
Wolf_China_Shanxi1
Tibetanwolf_BH4
China_SRR20326479_Sichuan
Wolf_Qinghai_CAN16
Tibetanwolf_MYL_SAMN12565073
Wolf_Tibetan_CAN32
Wolf_Tibetan_CAN9A
X
−0.00050
−0.00025
0.00000
0.00025
0.00050
D value
Zscore > -3
Zscore < -3

**Figure S11.** D statistic values to assess derived allele sharing between each wolf individual in Eurasia (X) with African wolves with the topology: (Graywolf$_{Norway}$, X), African wolf), Andeanfox). We used 4 African wolf genomes from Kenya, Morocco, Algeria, and Ethiopia as a population for P3. A negative D-value indicates an excess derived allele sharing with African wolves, while a positive D value can indicate allele sharing between X and the Andean fox, or X and the Gray wolf from Norway. Yellow color indicates D values that show statistically significant derived allele sharing with the Indian wolf (Z-score < -3). Wolves in purple indicate those that belong to the Indian lineage, greenish blue to Southwest Asia, and yellow are those that are from Pakistan.

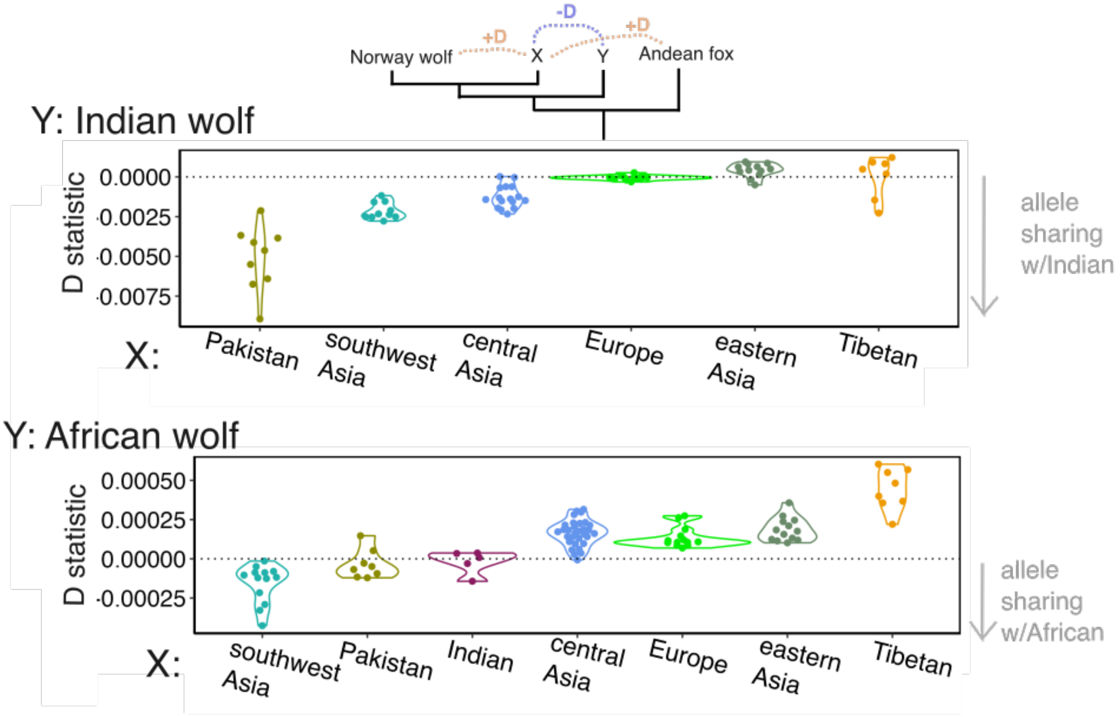


**Figure S12.** Derived allele sharing between each wolf individual by population with the Indian wolf and the African wolf. A negative D-value indicates an excess derived allele sharing with population Y, while a positive D value can indicate allele sharing between X and the Andean fox, or X and the gray wolf from Norway.

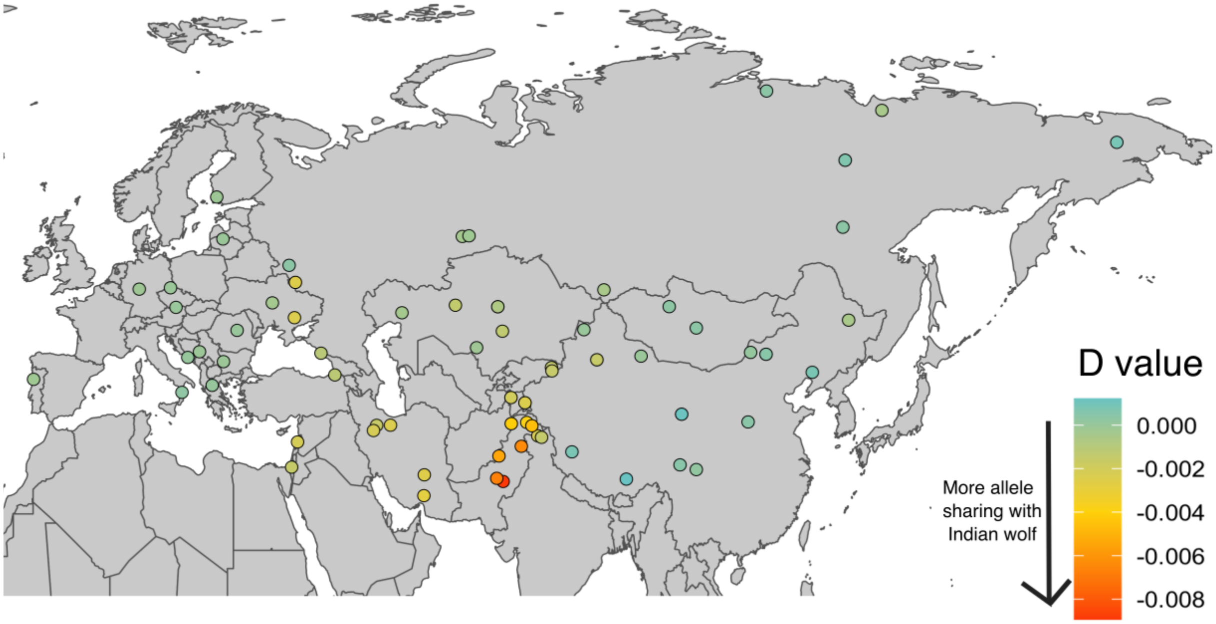


**Figure S13.** Derived allele sharing between each wolf individual in Eurasia (circles) with the Indian wolf with the topology: (Graywolf$_{Norway}$, X), Indian wolf–BH123), Andeanfox). A negative D-value indicates an excess derived allele sharing with the Indian wolf, while a positive D value can indicate allele sharing between X and the Andean fox, or X and the Gray wolf from Norway.

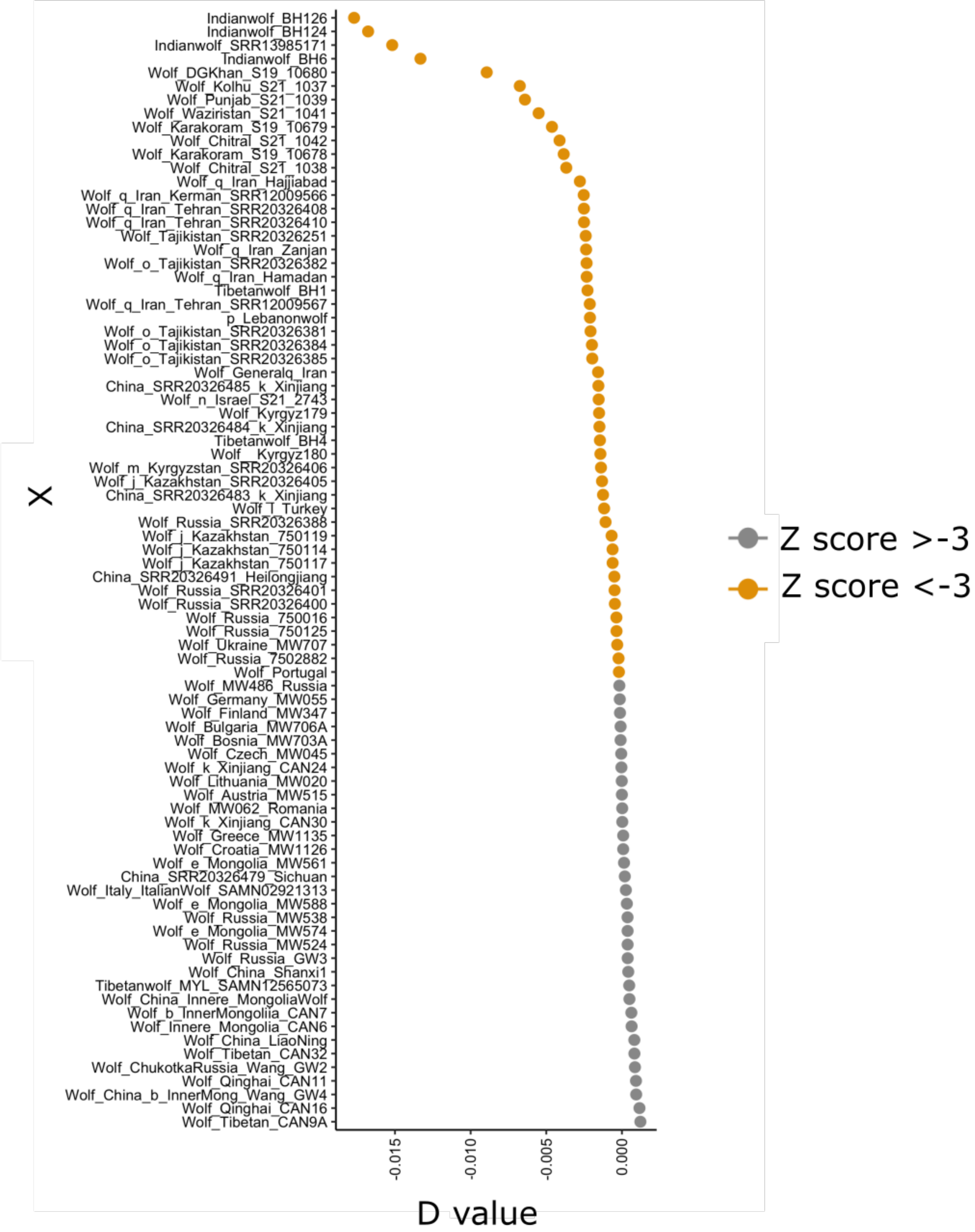


**Figure S14.** D statistic values to assess derived allele sharing between each wolf individual in Eurasia (X) with the Indian wolf with the topology: (Graywolf$_{Norway}$, X), Indian wolf BH123),

Andeanfox). A negative D-value indicates an excess derived allele sharing with the Indian wolf, while a positive D value can indicate allele sharing between X and the Andean fox, or X and the Gray wolf from Norway. Yellow color indicates D values that show statistically signifit derived allele sharing with the Indian wolf (Z-score < -3).

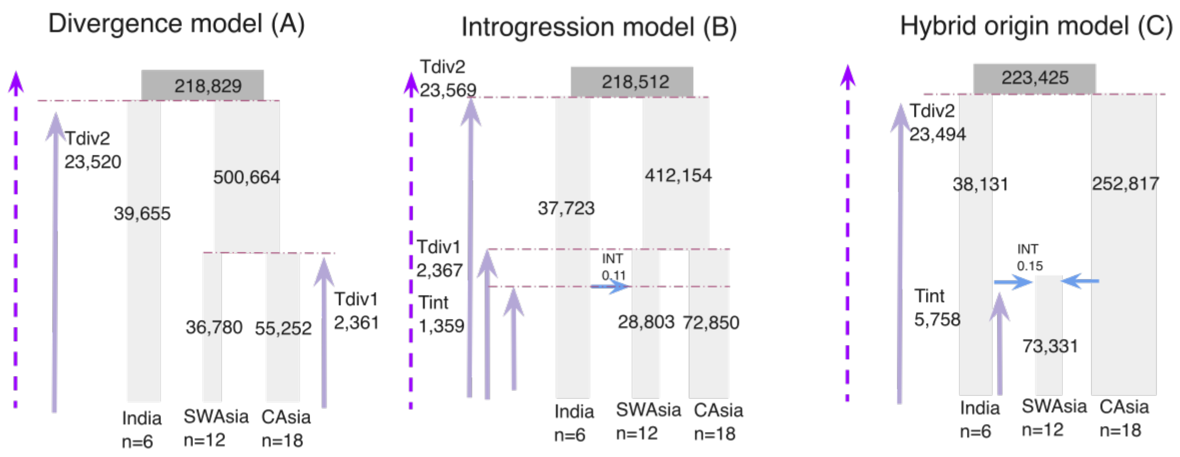


**Figure S15.** Three alternative scenarios for the origin of the Southwest Asian population tested in fastsimcoal2 (Excoffier et al. 2021). (A) A simple bifurcation model in which southwest Asia and central Asia diverged simultaneously from an ancestral Asian population; (B) an introgression model in which the present southwest Asian population was formed by introgression from India and central Asian wolves; (C) an hybridization model in which southwest Asia is an hybrid formed through admixture between wolves from India and central Asia. Note that model B is nested in model A, reducing A when INT=0.

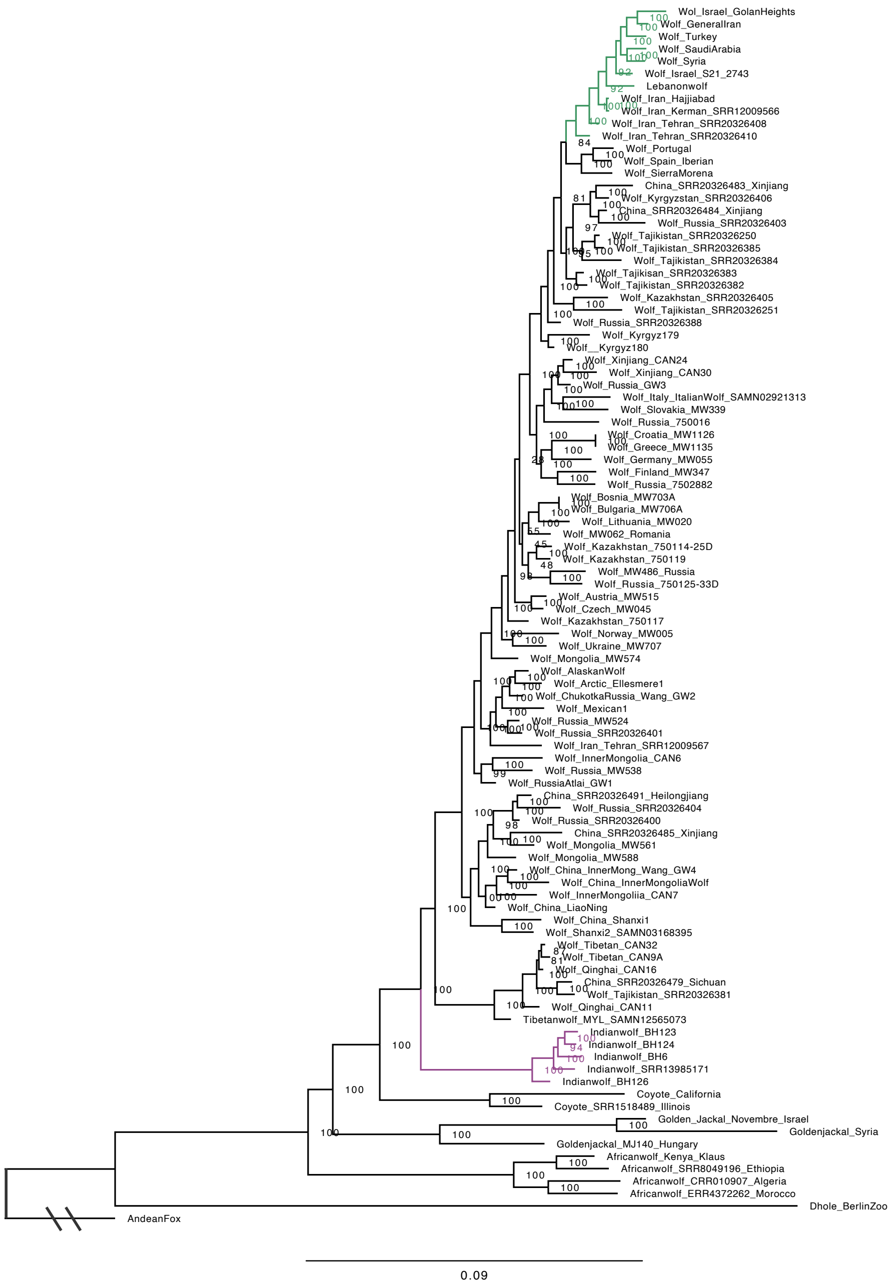


**Figure S16**. Fully labeled phylogenetic tree of the X chromosome using 514,048 SNPs found in only the low recombination regions (<0.2cM/Mb) with excluding wolves from Pakistan and two wolves from southwest Asia that were clustered with Indian wolves (wolf from Zanjan and Hamadan in Iran). We ran IQ-Tree 1.6.12 with estimating the best model using ModelFinder, which was TVM+F+R2, and used 1,000 ultra-fast bootstraps to infer each tree (Kalyaanamoorthy et al. 2017, Nguyen et al. 2014). The ultra-fast bootstrap support values are shown, where a 95% support corresponds to a probability of 95% the clade is correct.

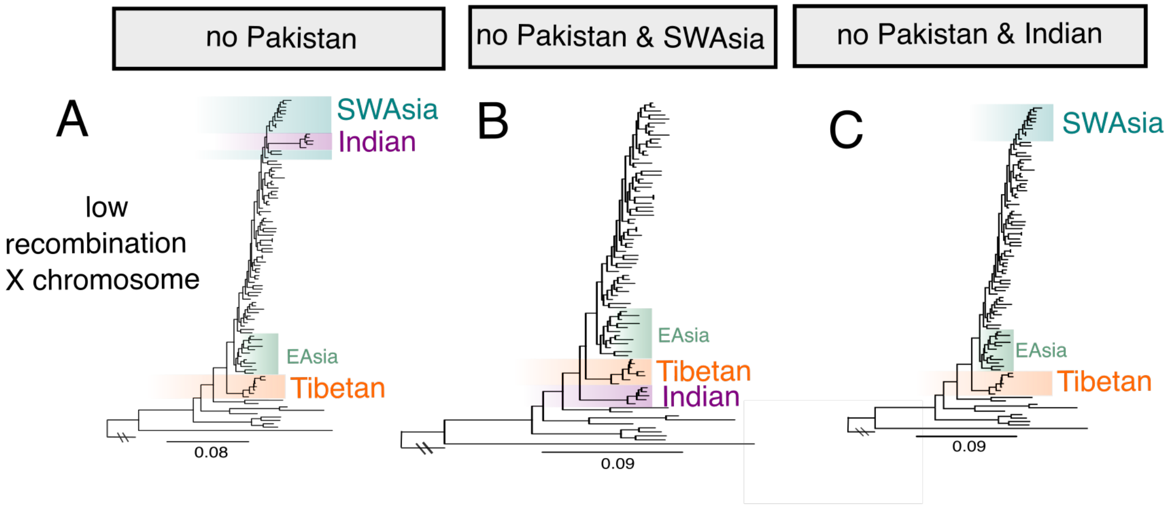


**Figure S17**. (A,B,C) Maximum likelihood autosomal phylogeny of wild canids inferred with IQ-Tree 1.6.12 using only low recombination (<0.2cM/Mb) regions of the X chromosome, we used IQ-Tree 1.6.12 where we estimated the best model using ModelFinder and used 1,000 ultra-fast bootstraps to infer each tree. Phylogenies vary in the wolf individuals they include: (A) has no wolves from Pakistan, (B) has no wolves from Pakistan and no wolves from southwestern (SW) Asia, and (C) has no wolves from Pakistan and no wolves from India. For tree A the best model was TVM+F+R2, tree B and C was TVM+F+R3.

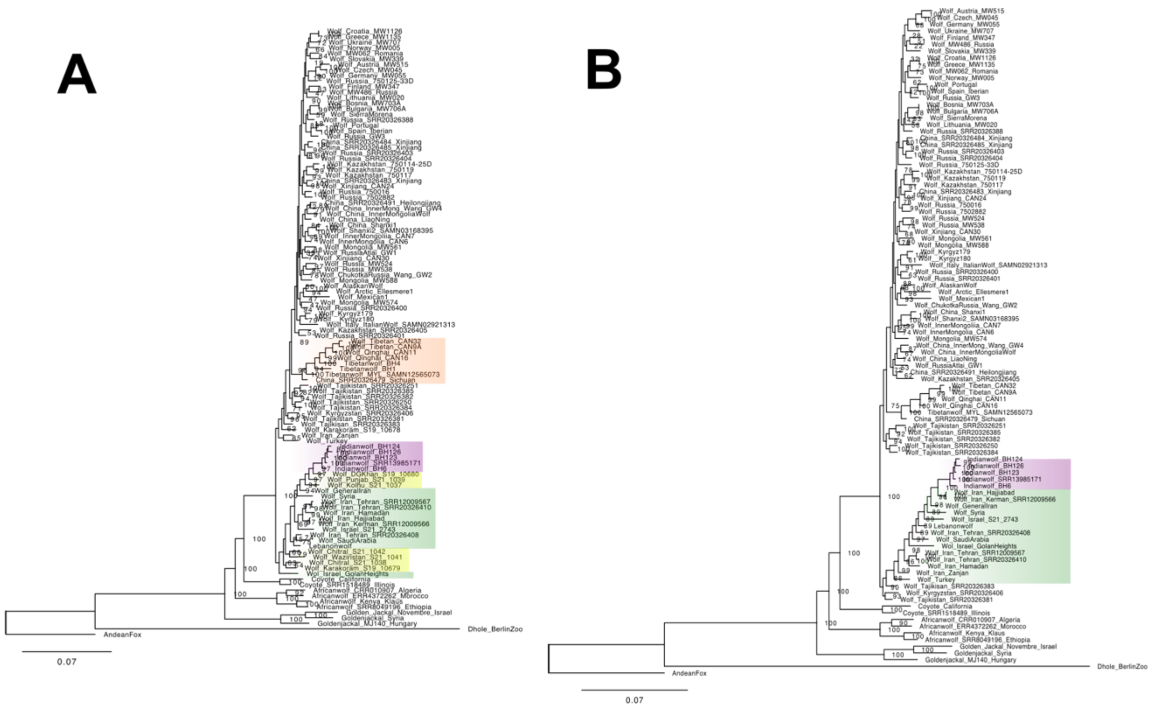


**Figure S18**. **(A)** Fully labeled maximum likelihood phylogeny using all wild canid individuals excepts dogs and wolves from Pakistan and Ladakh inferred using autosomal genomic regions in which the topology weight of topology 73 was above 0.3 (~99.99th percentile; ~40kb length total). The tree was inferred using IQ-tree where the best model was TVM+F+R4 through ModelFinder and 1000 bootstraps were used. Local posterior probability is labeled at selected nodes. **(B)** A maximum likelihood phylogeny with all wild canid individuals, which was inferred using autosomal genomic regions where the topology weight of topology 73 was above 0.3 (~99.99th percentile; ~40kb length total). The best model was TVM+F+R4 inferred using ModelFinder.

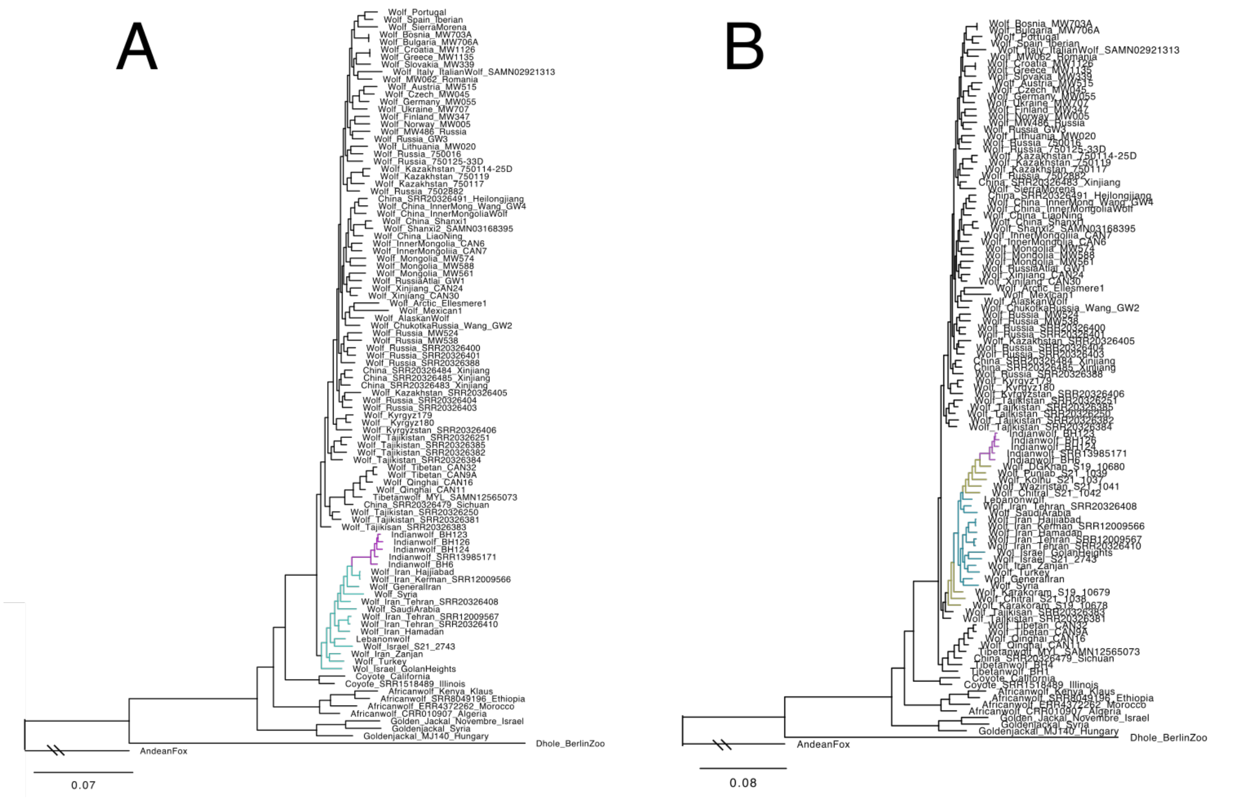


**Figure S19**. **(A)** A maximum likelihood phylogeny using all wild canid individuals excepts dogs and wolves from Pakistan and Ladakh inferred using autosomal genomic regions in which the topology weight of topology 73 was above 0.2 (~99.5$^{th}$ percentile; ~299kb length total). The tree was inferred using IQ-tree where the best model was xxx through ModelFinder and 1000 bootstraps were used. Local posterior probability is labeled at selected nodes. **(B)** A maximum likelihood phylogeny with all wild canid individuals, which was inferred using autosomal genomic regions where the topology weight of topology 73 was above 0.2 (~99.5$^{th}$ percentile; ~299kb length total).

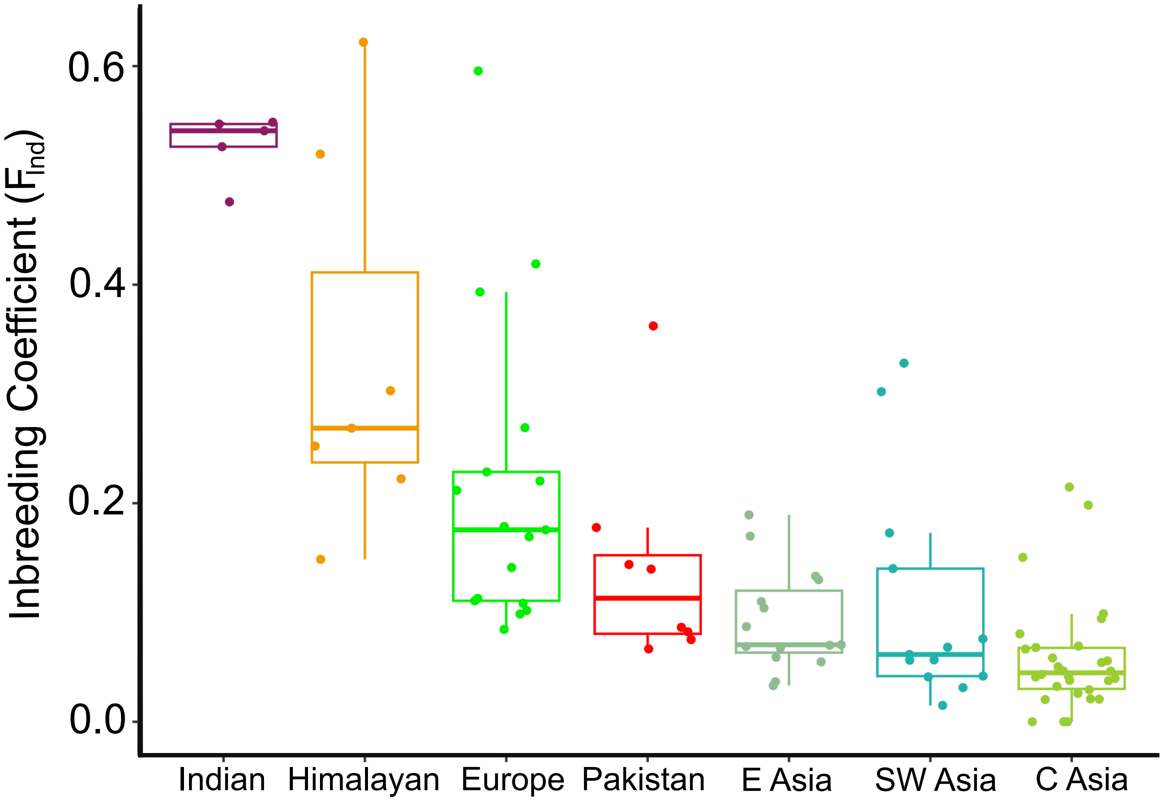


**Figure S20**. Estimated inbreeding coefficients ($F_{Ind}$) using genotype likelihoods with NgsRelate for 98 individuals across seven wolf populations in Eurasia.

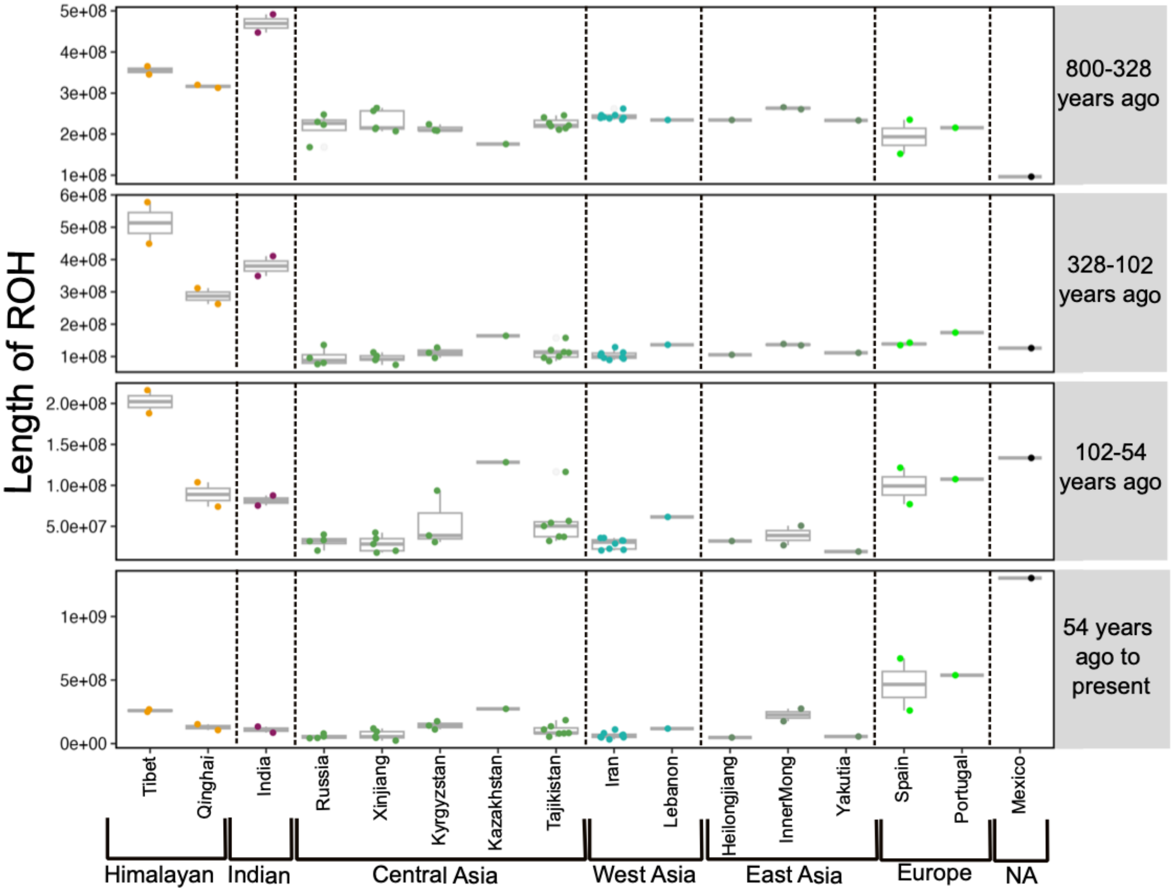


**Figure S21. Timing of inbreeding estimated from the length of ROH blocks across the autosomes of 42 wolves**. Inbreeding timings between 820-328 years ago corresponding to ROH lengths of 200Kb-500Kb, 328-102 years ago corresponding to ROH lengths of 500Kb-1.6Mb, 102-54 years ago corresponding to ROH lengths of 1.6Mb-3Mb, and 54 years ago to present corresponding to ROH lengths above 3Mb. Wolves most representing the Indian and Himalayan lineages generally show historical inbreeding in the last 800 years compared to wolves from Mexico and Spain, which show recent inbreeding in the 100 years.

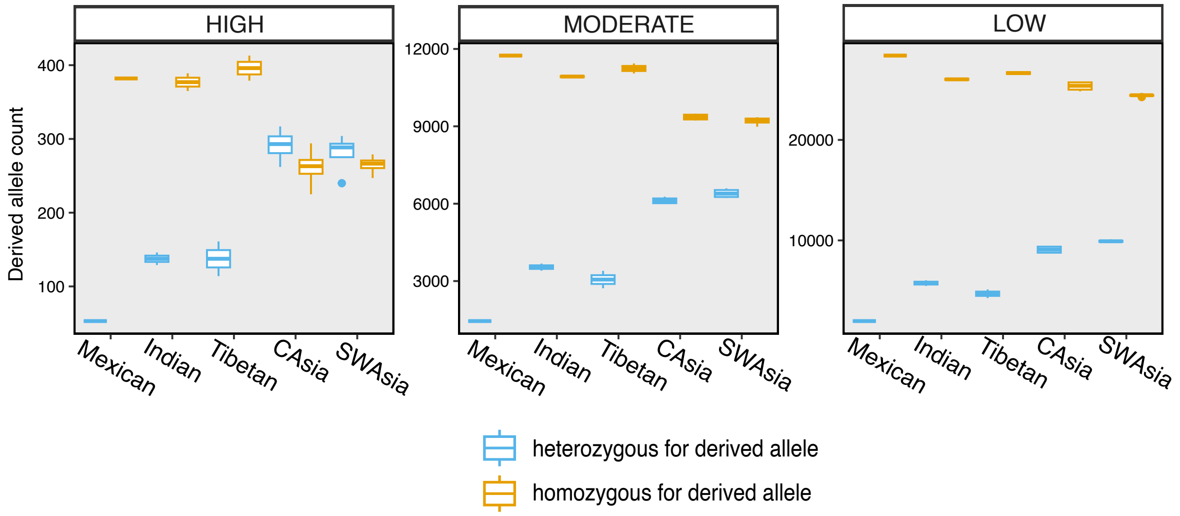


**Figure S22.** Number of heterozygous and homozygous counts for the derived allele for High impact, Medium impact, and Low impact categories. Homozygous counts contain two derived alleles and heterozygous derived counts contain one derived allele. For High and Moderate categories, we find Mexican, Indian, and Tibetan wolves are hgher counts of homozygous derived alleles, consistent with genome-wide measures of higher homozygosity.

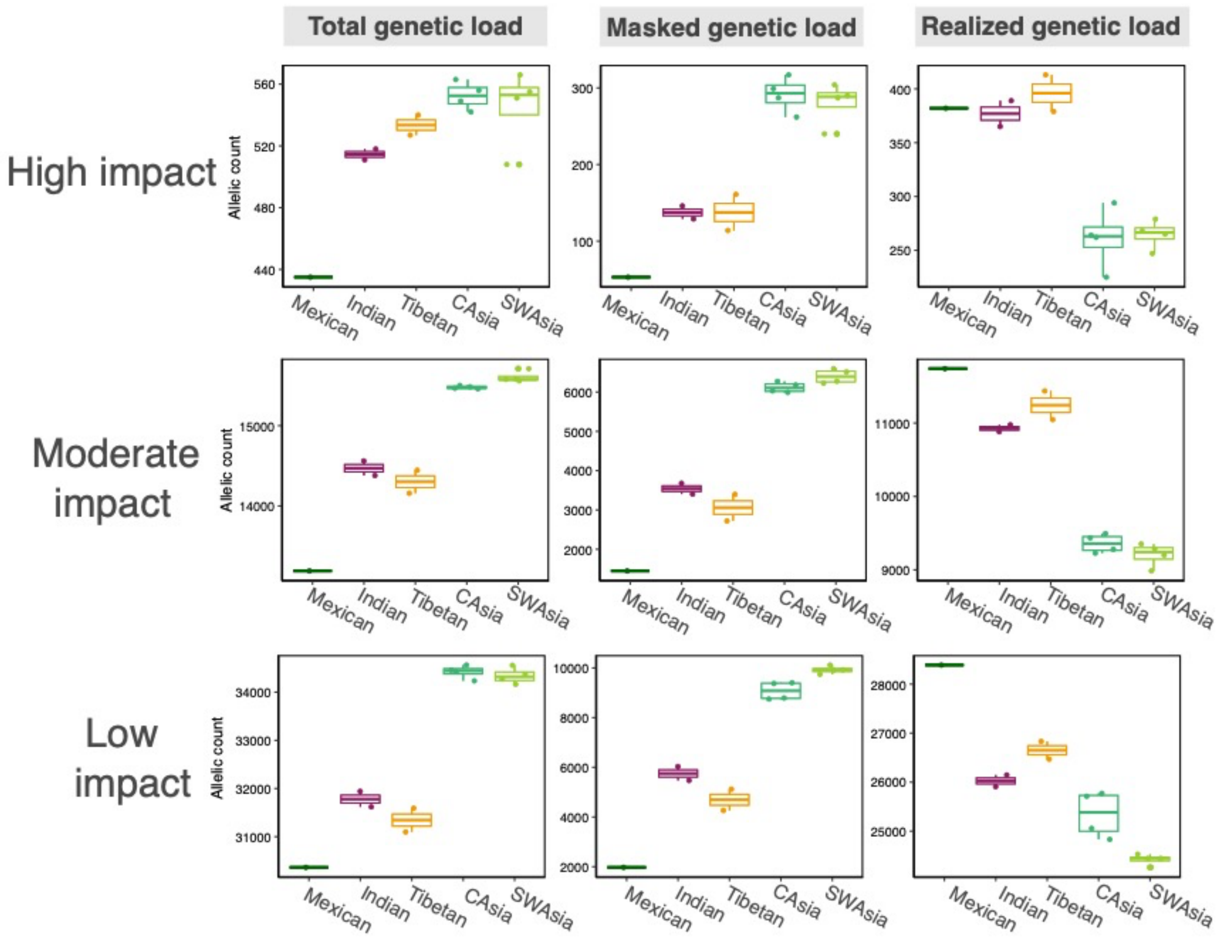


**Figure S23.** Total genetic load (the total number of derived alleles; homozygous derived alleles counted twice, heterozygous derived alleles once), realized load (homozygous state of derived alleles), and masked load (heterozygous state of derived alleles) for each impact category (High, Moderate, Low) for five selected wolf populations of wolves. For all categories, we find Mexican, Indian, and Tibetan wolves have a lower total genetic load than wolves in southwest Asia and central Asia. Because low impact is considered to be mostly harmless, the lower total genetic load in historically small wolf populations suggests it could be due to loss of total derived variants from genetic drift.

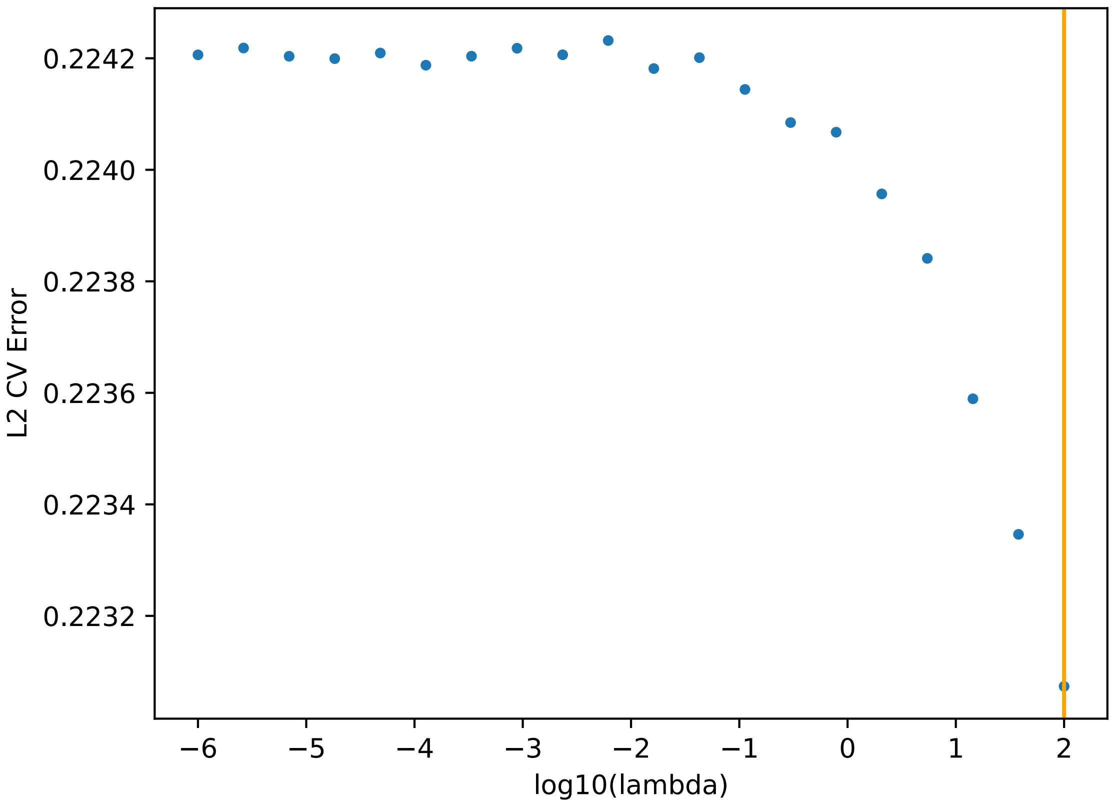


**Figure S24**. Cross validation results by using leave-one-out cross validation to select the optimal value of lambda, the smoothing parameter. A lambda of 100 was selected to be the optimal value and used in the final analysis.

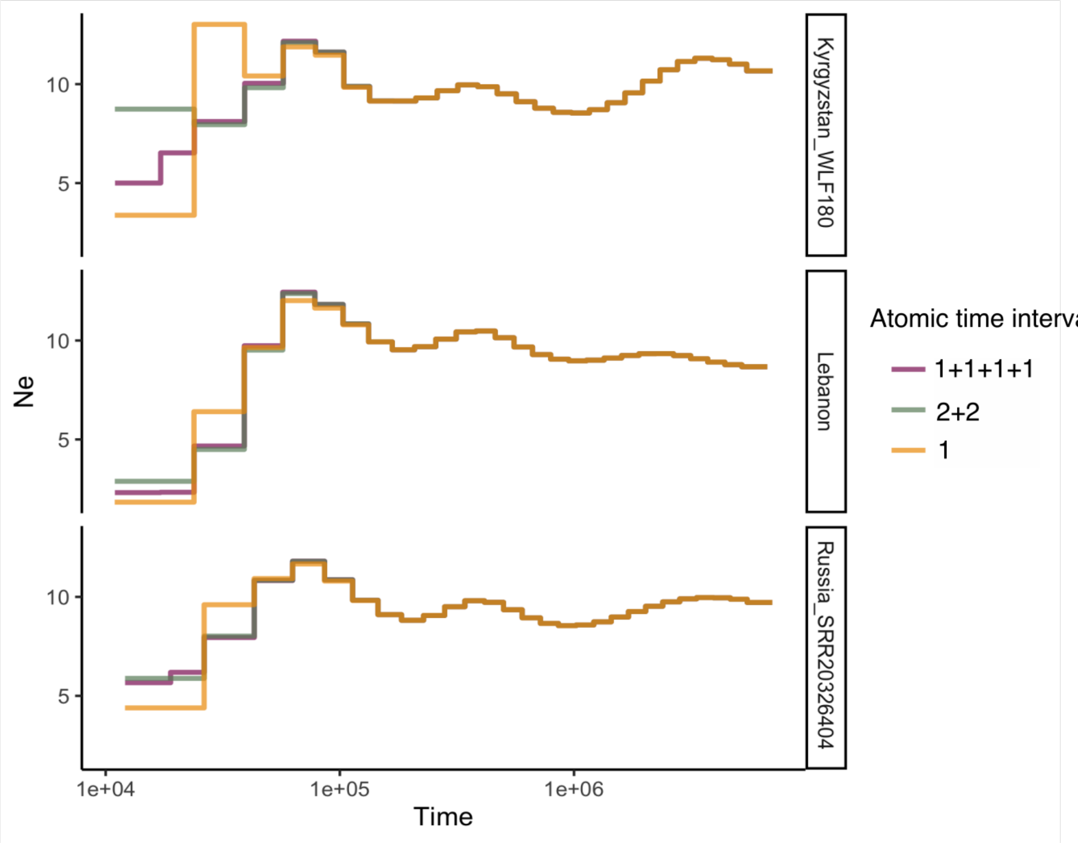


**Figure S25.** PSMC plots for three wolf genomes while varying the atomic time intervals. We observe erroneous peaks for the Kyrgyzstan WLF180 genome when using atomic time interval 1, whereas this peak disappears when using 2+2 or 1+1+1+1. In our study, we use the atomic time interval of 1+1+1+1 to avoid false PSMC peaks.

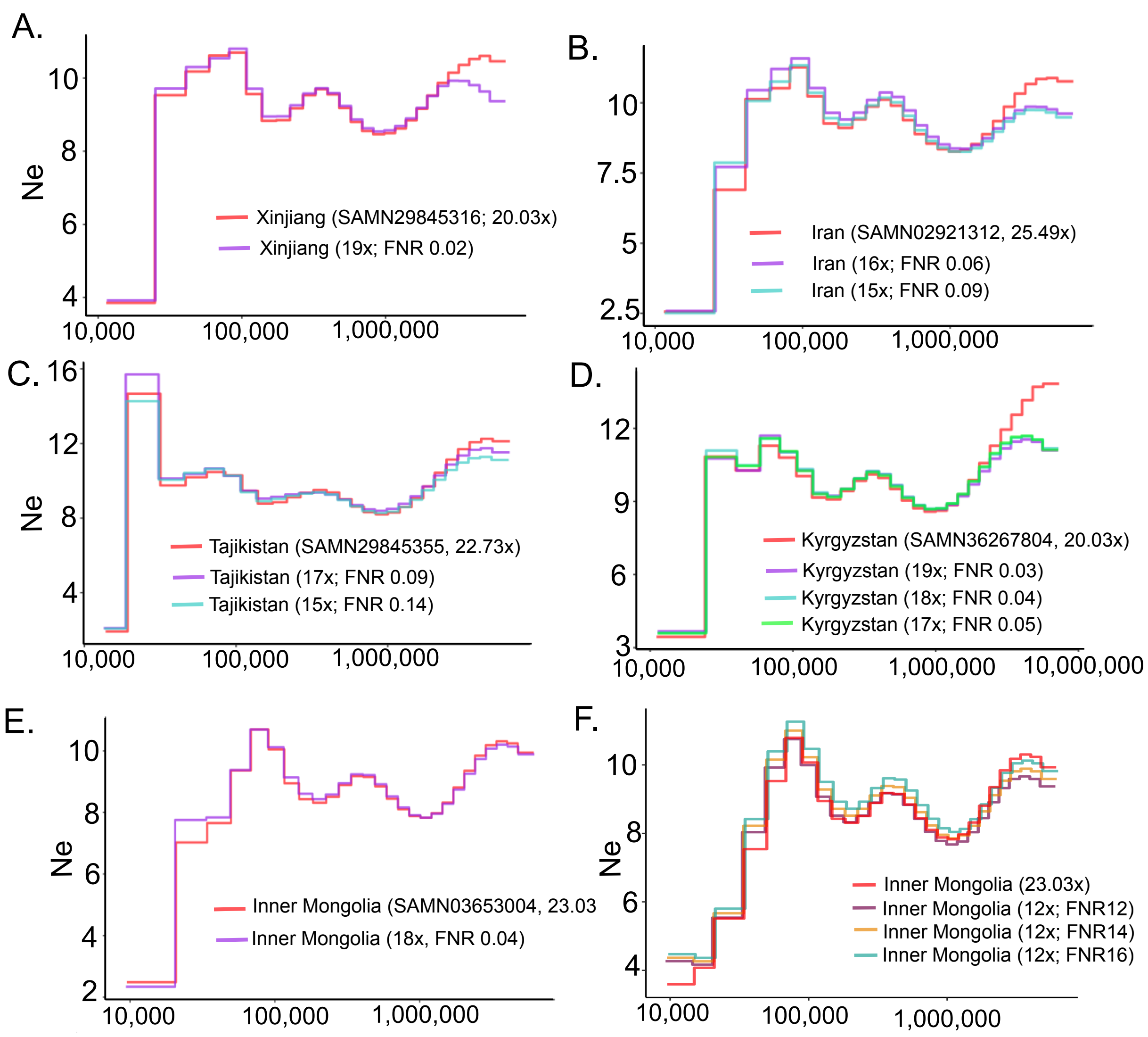


**Figure S26**. Down-sampling of genomes to select the false negative rates (FNR) for low-depth corrections of pairwise sequentially Markovian coalescent (PSMC) demographic trajectories. High coverage wolf genomes (>20x) that were from the same region as the target low coverage samples were selected, which consisted of genomes from Xinjiang (SAMN29845316; 20.03x), Iran (SAMN02921312, 25.49x), Inner Mongolia (SAMN03653004, 23.03x), Kyrgyzstan (SAMN36267804, 25.86), and Tajikistan (SAMN29845355, 22.73x). Panel F was used to select the false negative rate (FNR) for the wolf genome from Shanxi, China, which was at a coverage of 12x.

**Figure S27.** Est and Tpl input files for fastsimcoal2 for each of the three models we tested.

**Model 1: 3PopDiv**

**Description**: 3PopDiv ('Strictly biforcating'): One single population diverged into two pops (India and Asia) at a given time TSPLIT in the past, and after that, at time TDIV1, there was a split of the 'Asia' population into two populations (CAsia and SWAsia) so that at present time there is three populations: India SWAsia and CAsia

**3PopDiv.est**

---

```
// Search ranges and rules file
// ****************************

[PARAMETERS]
//#isInt? #name   #dist.#min  #max
//all Ns are in number of haploid individuals
1  $NCAS$      unif  2e3   4e5   output
1  $NIND$      unif  2e3   4e5   output
1  $NSWAS$      unif  2e3   4e5   output
1  $NASANC$      unif 2e3   4e5   output
1  $NANC$      unif  2e3   4e5   output
```

```
1  $TSPLIT$   unif  20000   50000  output bounded
1  $TDIV1$      unif  2325   $TSPLIT$     output bounded paramInRange

[COMPLEX PARAMETERS]
```

---

**3PopDiv.tpl**

---

```
//Parameters for the coalescence simulation program : fsimcoal2.exe
3 samples to simulate :
//Population effective sizes (number of genes)
$NCAS$ //cAsia
$NIND$                                  //India
$NSWAS$                                 //swAsia
//Samples sizes and samples age
18
6
12
//Growth rates                          : negative growth implies population expansion
0
0
0
//Number of migration matrices : 0 implies no migration between demes
0
//historical event: time, source, sink, migrants, new deme size, new growth rate, migration matrix index
2 historical event
$TDIV1$ 0 2 1 $NASANC$ 0 0 absoluteResize
$TSPLIT$ 2 1 1 $NANC$ 0 0 absoluteResize
//Number of independent loci [chromosome]
1 0
//Per chromosome: Number of contiguous linkage Block: a block is a set of contiguous loci
1
//per Block:data type, number of loci, per generation recombination and mutation rates and optional parameters
FREQ  1   0  4.5e-9 OUTEXP
```

---

**<u>Model 2: </u>3PopIntrog**

**Description**: 3PopIntrog ('Introgression Model'): One single population diverged into two pops (India and Asia) at a given time TSPLIT in the past, and after that, at time TDIV1, there was a split of the 'Asia' population into two populations (CAsia and SWAsia) so that at present time there is India SWAsia and CAsia. At a given time TINTROG in the past there was introgression from India to SWAsia (proportion INT)

**3PopIntrog.est**

---

```
// Search ranges and rules file
// ****************************
```

```
[PARAMETERS]
//#isInt? #name   #dist.#min  #max
//all Ns are in number of haploid individuals
1  $NCAS$      unif  2e3   4e5  output
1  $NIND$      unif  2e3   4e5   output
1  $NSWAS$      unif  2e3   4e5   output
1  $NASANC$      unif 2e3   4e5   output
1  $NANC$      unif  2e3   4e5   output

1  $TSPLIT$   unif  20000    50000  output bounded
1  $TDIV1$      unif  2325   $TSPLIT$     output bounded paramInRange
1  $TINTROG$       unif  1   $TDIV$    output bounded paramInRange

0  $INT$    logunif  0.000000001 1 output bounded

[COMPLEX PARAMETERS]
```

---

**3PopIntrog.tpl**

---

```
//Parameters for the coalescence simulation program : fsimcoal2.exe
3 samples to simulate :
//Population effective sizes (number of genes)
$NCAS$ //cAsia
$NIND$ //India
$NSWAS$ //swAsia
//Samples sizes and samples age
18
6
12
//Growth rates                                    : negative growth implies population expansion
0
0
0
//Number of migration matrices : 0 implies no migration between demes
0
//historical event: time, source, sink, migrants, new deme size, new growth rate, migration matrix index
3 historical event
$TINTROG$ 2 1 $INT$ 1 0 0
$TDIV$ 0 2 1 $NASANC$ 0 0 absoluteResize
$TSPLIT$ 2 1 1 $NANC$ 0 0 absoluteResize
//Number of independent loci [chromosome]
1 0
//Per chromosome: Number of contiguous linkage Block: a block is a set of contiguous loci
1
//per Block:data type, number of loci, per generation recombination and mutation rates and optional
parameters
FREQ  1   0  4.5e-9 OUTEXP
```

---

**Model 3: 3PopHyb**
**Description:** 3PopHyb ('Hybrid origin'): One single pop diverged into two pops India and CAsia at a given time TSPLIT in the past, and after that at time TINTROG a new hybrid population (SWasia) was formed by the contribution of INT from India and (1-INT) from CAsia.

**3PopHyb.est**

---

```
// Search ranges and rules file
// ****************************

[PARAMETERS]
//#isInt? #name   #dist.#min  #max
//all Ns are in number of haploid individuals
1  $NCAS$      unif  2e3   4e5   output
1  $NIND$      unif  2e3   4e5   output
1  $NSWAS$      unif  2e3   4e5   output
1  $NANC$      unif  2e3   4e5   output

1  $TSPLIT$   unif  20000    50000  output bounded
1  $TINTROG$      unif  1   $TSPLIT$    output bounded paramInRange

0  $INT$    logunif  0.000000001 1 output bounded

[COMPLEX PARAMETERS]
1 $TINTROG20$ = $TINTROG$+1 hide
```

---

**3PopHyb.tpl**

---

```
//Parameters for the coalescence simulation program : fsimcoal2.exe
3 samples to simulate :
//Population effective sizes (number of genes)
$NCAS$ //cAsia
$NIND$                                    //India
$NSWAS$                                   //swAsia
//Samples sizes and samples age
18
6
12
//Growth rates                            : negative growth implies population expansion
0
0
0
//Number of migration matrices : 0 implies no migration between demes
```

```
0
//historical event: time, source, sink, migrants, new deme size, new growth rate, migration matrix index
3 historical event
$TINTROG$ 2 1 $INT$ 1 0 0
$TINTROG20$ 2 0 1 1 0 0
$TSPLIT$ 1 0 1 $NANC$ 0 0 absoluteResize
//Number of independent loci [chromosome]
1 0
//Per chromosome: Number of contiguous linkage Block: a block is a set of contiguous loci
1
//per Block:data type, number of loci, per generation recombination and mutation rates and optional parameters
FREQ  1   0  4.5e-9 OUTEXP
```

---
